# A reaction-diffusion framework for *de novo* Polycomb spreading

**DOI:** 10.64898/2026.08.26.747395

**Authors:** Eleanor A. Degen, Shelby A. Blythe

## Abstract

Eukaryotic organisms rely on post-translational modifications to chromatin to maintain stable patterns of gene silencing. These modifications include trimethylation at histone H3 lysine 27 (H3K27me3), which is deposited by Polycomb Repressive Complex 2 (PRC2) and accumulates on the genome during embryogenesis. While this process underlies the proper specification of cell types, we lack the ability to quantitatively predict the *de novo* establishment of Polycomb states. The kinetics of H3K27 methylation is difficult to quantify *in vivo*, and further, the network of molecular interactions that influences Polycomb states is complex. Here, leveraging the *Drosophila* embryonic system, we measure H3K27me3 dynamics with ChIP-seq and extract the rate the modification spreads along chromatin *in vivo*. To provide a mechanistic explanation for this rate, we build a reaction-diffusion framework that models how PRC2 establishes states of gene silencing *de novo*. The reaction-diffusion system recapitulates experimental observations in wild-type and mutant embryos, and suggests that PRC2 can diffuse in 1D along chromatin at a rate enhanced by Polycomb Repressive Complex 1. Through this work, we define a minimal set of parameters that dictate *in vivo* Polycomb dynamics, and provide evidence that the early embryo creates a “super-charged” environment for epigenetic modification.

## INTRODUCTION

Eukaryotic genomes are wrapped around nucleosomes to form chromatin fibers that fit within cells. How cells regulate gene expression depends on this packaging. Chemical modifications on the histone subunits of nucleosomes can facilitate or silence gene expression (Millán-Zambrano et al., 2022; Strahl & Allis, 2000). Across species, Polycomb Group (PcG) proteins establish and maintain histone modifications associated with inactive gene expression states (Grossniklaus & Paro, 2014; Laugesen et al., 2019; Margueron & Reinberg, 2011; Schuettengruber et al., 2017). PcG proteins are critical for gene silencing over the lifespan of the organism, yet quantitatively predicting Polycomb modification dynamics is challenging, as it requires that we first understand both PcG protein localization on chromatin and the *in vivo* catalytic activity of the system. Computational modeling can help decode the mechanisms that drive the establishment of silent chromatin states.

To benchmark a model, one must be able to measure Polycomb modifications as they spread across the genome *de novo*, a process challenging to observe in mature cells that maintain a stable chromatin landscape. In the cells of adult organisms or in late embryonic contexts, the primary disruption to the epigenetic state occurs following cell division, when unmodified histones are incorporated into chromatin during genome replication (Alabert & Groth, 2012; Groth et al., 2007). To maintain Polycomb states, Polycomb Repressive Complex 2 (PRC2) is recruited to sites of existing Polycomb modifications to catalyze the canonical mark of Polycomb silencing, trimethylation at histone H3 lysine 27 (H3K27me3), as well as lower-order H3K27 methylation states (H3K27me1/2) (Hansen et al., 2008; Margueron et al., 2009). Despite substantial modeling efforts to describe the stable maintenance of H3K27me3 across cell divisions (Lövkvist & Howard, 2021; Lundkvist et al., 2023; Owen et al., 2023), no prior models have fully addressed how PRC2 spreads H3K27me3 along chromatin *de novo*, a process that shapes the chromatin landscape during development. The establishment of H3K27me3 in early embryos of the fruit fly *Drosophila melanogaster* has nonetheless been measured (Gonzaga-Saavedra et al., 2026; Li et al., 2014; Reinig et al., 2020). These out-of-equilibrium measurements can help guide the building of a mathematical framework that addresses outstanding questions about the Polycomb system.

Studies have measured the kinetics of the catalytic subunit of PRC2, Enhancer of Zeste (E(z), flies) / Enhancer of Zeste Homolog 1/2 (EZH1/2, mammals) (Cao et al., 2002; Czermin et al., 2002; McCabe et al., 2012; Müller et al., 2002; Sneeringer et al., 2010; Y. Zheng et al., 2012), yet we do not know how kinetic parameters translate to the spreading of Polycomb modifications along chromatin. The effective catalytic rate constants for H3K27 methylation reactions measured in cell culture are on the order of day^-1^ (Y. Zheng et al., 2012), contrasting the observation that detectable H3K27me3 accumulates over a ∼1 hour cell cycle in the *Drosophila* embryo (Gonzaga-Saavedra et al., 2026; Li et al., 2014). Further, computational modeling suggests that the probability that E(z) methylates histones is an order of magnitude higher in *Drosophila* embryos than cultured *Drosophila* cells (Lundkvist et al., 2023). The early embryo may therefore represent a remarkably enhanced environment for the establishment of Polycomb states. While the *Drosophila* embryo establishes H3K27me3 over an hour, in mammalian cell lines, genetic and pharmacological methods for measuring kinetics have revealed H3K27me3 accumulates on chromatin over days (Højfeldt et al., 2018; Oksuz et al., 2018; Reverón-Gómez et al., 2018; Veronezi & Ramachandran, 2024). Low levels of H3K27me3 are detectable at a small subset of loci after 4 hours of establishment in mouse embryonic stem cells, but the modification takes at least four days to reach an equilibrium (Højfeldt et al., 2018; Veronezi & Ramachandran, 2024). In addition to flies, H3K27me3 kinetics in mouse, fish, and frog embryos likely outpace those of mammalian cell culture. In early mouse development, the paternal X chromosome gains H3K27me3 sharply between the morula and blastocyst stages, within a 12-24 hour period (Matsuwaka et al., 2025). In zebrafish, H3K27me3 domains are established shortly after zygotic genome activation (∼4 hours post-fertilization) (Hickey et al., 2022). In *Xenopus* development, high levels of H3K27me3 accumulate on chromatin in the ∼5 hours spanning stage 9 and stage 12 (Akkers et al., 2009; van Heeringen et al., 2014). However, while early embryonic Polycomb states have been measured in mouse, fish, and frog, the low frequency of experimental sampling in these studies limits our ability to make quantitative conclusions (Akkers et al., 2009; Hickey et al., 2022; Matsuwaka et al., 2025; Mei et al., 2021; van Heeringen et al., 2014; H. Zheng et al., 2016). How molecular parameters define the spreading of Polycomb modifications along chromatin during embryogenesis remains an open question.

Here, we quantify the rate that H3K27me3 spreads over chromatin in the *Drosophila* embryo, and develop a reaction-diffusion framework that recapitulates these dynamics. The model provides evidence that PRC2 progresses along the genome effectively according to a 1-dimensional random walk, at a rate approximately 10x slower than the reported sliding of transcription factors along DNA. By applying the reaction-diffusion model to predict the results of experimental perturbation to the Polycomb system, we demonstrate that H2A ubiquitylation associated with the Sce/dRing1 subunit of Polycomb Repressive Complex 1 substantially increases the speed of PRC2 progression along chromatin and the rate of H3K27me3 spreading. We further define the specialized Polycomb conditions of embryogenesis by evaluating how altering the model allows for predicting H3K27me3 accumulation in mammalian cell culture. Through this work, we provide evidence that E(z)/EZH2 operates with effectively elevated catalytic efficiencies in early embryos, allowing for rapid remodeling of the Polycomb landscape during development.

## RESULTS

### Quantification of *in vivo* H3K27me3 spreading rates

We drew upon the *Drosophila* embryo system to quantify how H3K27me3 spreads out from sites of E(z) nucleation on chromatin, Polycomb Response Elements (PREs). We extended our prior time course of chromatin immunoprecipitation sequencing (ChIP-seq) measurements of H3K27me3 over the ∼1-hour cell cycle following mitosis 13 of *Drosophila* development (Nuclear Cycle 14, NC14) (Gonzaga-Saavedra et al., 2026) by measuring H3K27me3 in stage 9 (NC14 + 2h), stage 11 (NC14 + 4h), and stage 12 (NC14 + 6h) embryos. These measurements indicate that, after H3K27me3 appears local to PREs at mid-NC14 (NC14 + 35’) (Gonzaga-Saavedra et al., 2026), the modification continually spreads out from these sites over the next ∼6 hours (Fig. 1A & B). The *de novo* establishment of H3K27me3 following mitosis 13 through a nucleation and spreading mechanism provides a platform for a quantitative analysis of H3K27me3 spreading rates.

**Figure 1.**
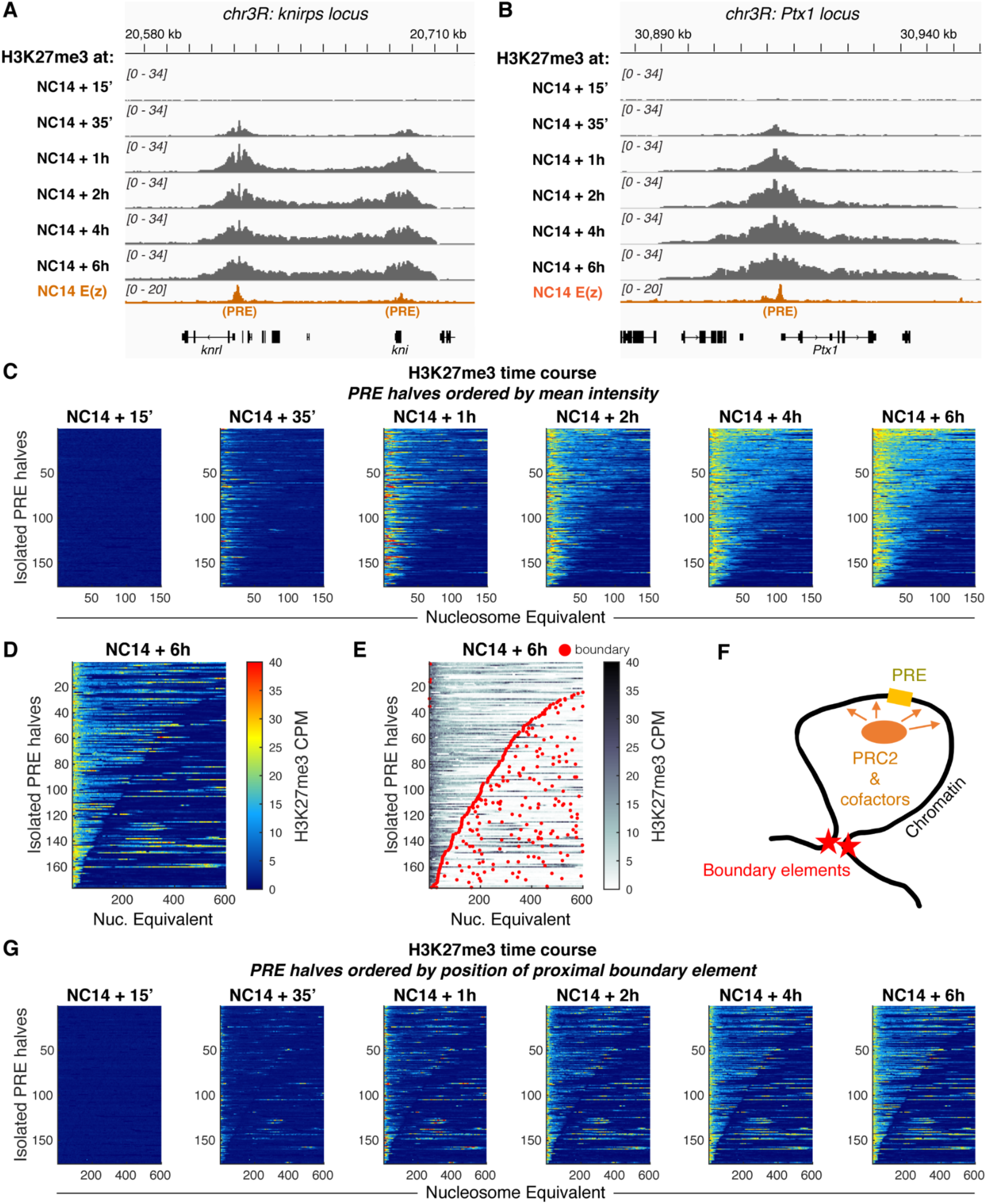
Analysis of H3K27me3 dynamics given higher-order chromatin structure. **A)** H3K27me3 and E(z) occupancy at the *knirps* locus. The top three rows show counts per million (CPM) normalized ChIP-seq measurements of H3K27me3 at early-NC14 (NC14 + 15’), mid-NC14 (NC14 + 35’), and late-NC14 (NC14 + 1h) time points, data replotted from Gonzaga-Saavedra et al. (Gonzaga-Saavedra et al., 2026). Rows 4-6 show CPM-normalized ChIP-seq measurements of H3K27me3 at embryonic stage 9 (NC14 + 2h), stage 11 (NC14 + 4h), and stage 12 (NC14 + 6h), newly collected data in this work. The bottom row shows CPM-normalized ChIP-seq measurements of E(z) in NC14 embryos, data replotted from Gonzaga-Saavedra et al. (Gonzaga-Saavedra et al., 2026). E(z) peaks mark PREs. H3K27me3 spreads out bidirectionally from these PREs over six hours following mitosis 13. **B)** H3K27me3 and E(z) occupancy at the *Ptx1* locus. Rows show H3K27me3 and E(z) ChIP-seq data as defined in (A). **C)** H3K27me3 at isolated PRE halves over six hours following mitosis 13. Isolated PRE halves are defined in the Materials and Methods. In each heatmap, rows show mean CPM-normalized ChIP-seq measurements of H3K27me3 in 180 base pair bins, corresponding to “nucleosome equivalents.” Rows are ordered by mean intensity across 150 nucleosomes at NC14 + 6h, and are aligned by the position of the center of the E(z) peak in each PRE, which in the heatmaps is at nucleosome equivalent = 1. x-axes span 150 nucleosomes. Each heatmap has an equivalent color axis, showing a range of [0, 40] CPM-normalized ChIP-seq reads. NC14 + 15’, NC14 + 35’, and NC14 + 1h heatmaps present a re-analysis of data collected in Gonzaga-Saavedra et al. (Gonzaga-Saavedra et al., 2026). **D)** H3K27me3 at isolated PRE halves at NC14 + 6h ordered by the position of the first boundary element proximal to nucleosome equivalent = 0, the position of the E(z) peaks. x-axis spans 600 nucleosomes. The heatmap shows mean CPM-normalized ChIP-seq measurements of H3K27me3 in NC14 + 6h embryos. **E)** H3K27me3 at isolated PRE halves at NC14 + 6h ordered by the position of the first boundary element proximal to nucleosome equivalent = 0, overlaid with boundary element positions (red points) measured in Batut et al. (Batut et al., 2022). As in (D), the heatmap shows mean CPM-normalized ChIP-seq measurements of H3K27me3 in NC14+ 6h embryos. Boundary elements frequently mark the boundaries of H3K27me3 domains. **F)** A schematic of PRC2 (orange) activity at a PRE (yellow) within a TAD bounded by boundary elements (red). **G)** H3K27me3 spreads out from isolated PREs over six hours post-mitosis 13 until reaching flanking boundary elements. Heatmaps show mean CPM-normalized ChIP-seq measurements of H3K27me3 in NC14 + 15’, NC14 + 35’, NC14 + 1h, NC14 + 2h, NC14 + 4h, and NC14 + 6h embryos. In each heatmap, rows are ordered by the position of the first boundary element proximal to nucleosome equivalent = 0. As in (C), NC14 + 15’, NC14 + 35’, and NC14 + 1h heatmaps show an analysis of data collected in Gonzaga-Saavedra et al. (Gonzaga-Saavedra et al., 2026). x-axes span 600 nucleosomes.

To assess Polycomb activity across loci, for each time point we calculated the average H3K27me3 signal surrounding “isolated PREs” in the genome, E(z) nucleation sites that are ≥ 150 nucleosome equivalents away from their neighbors (Materials and Methods) (Degen et al., 2026). A “nucleosome equivalent” corresponds to 180 bp (approximately the footprint of a nucleosome + linker), and we refer to them as “nucleosomes” for convenience. We then evaluated how H3K27me3 spreads from these PREs in both the 5’ and 3’ directions by splitting each Polycomb domain at the position of their PRE, flipping the upstream segment, and aligning the domain halves by PRE position (Fig. 1C). Early in NC14 (NC14 + 15’), H3K27me3 is undetectable on chromatin (Fig. 1C). By NC14 + 35’, the modification has begun to accumulate close to the PREs, but has yet to spread farther out (Fig. 1C). Over the next ∼6 hours, at most loci H3K27me3 spreads over at least 150 nucleosomes (Fig. 1C). However, at a subset of PREs, H3K27me3 fails to progress past even 50 nucleosomes (Fig. 1C).

Boundary elements that partition the genome into topologically associating domains (TADs) could prevent PRC2 from methylating outside of the TADs containing its nucleation sites. Insulators, along with active histone marks and transcription units, can restrain the deposition of Polycomb modifications (De et al., 2019, 2020; Fujioka et al., 2013; Park et al., 2012; Schmitges et al., 2011; Yuan et al., 2011). We therefore sought to take into account the physical limits of Polycomb domains in order to define regions for analysis where H3K27me3 spreads freely. To designate regions of PRC2 spreading, we drew upon boundary positions determined previously by MicroC in NC14 *Drosophila* embryos (Batut et al., 2022), and ordered each H3K27me3 track at NC14 + 6h by the position of the first boundary element relative to each associated PRE (Fig. 1D). Annotating the ordered NC14 + 6h H3K27me3 measurements with boundary element positions highlights that these elements often mark the limits of Polycomb domains (Fig. 1E). Outside of the first boundary proximal to each isolated PRE, chromatin sometimes contains H3K27me3, but these patterns likely result from the presence of additional PRC2 nucleation sites ≥ 150 nucleosomes away from the isolated PREs (Fig. 1E). Many of the isolated PREs likely occupy TADs where boundary elements limit the extent to which PRC2 can spread H3K27me3 along chromatin (Fig. 1F). Over the six hours following mitosis 13, from most isolated PREs, we find H3K27me3 spreads until it reaches a boundary element (Fig. 1G). This observation supports prior observations that boundary elements limit PRC2 progression along the genome (De et al., 2019, 2020; Fujioka et al., 2013; Park et al., 2012), and allowed us to operationally define genomic regions for analysis where H3K27me3 spreading occurs.

Our measurements uniquely positioned us to calculate the rate that H3K27me3 spreads freely from PRC2 nucleation sites. To extract a spreading rate, we performed linear regression to fit exponentials to the H3K27me3 profiles while omitting any genomic regions outside of the nearest adjacent boundary elements (Fig. 2A). Omitting these regions, which are unlikely to be mechanistically linked to an individual PRE, helped us prevent incorrect association of external signal with a source of H3K27me3 spreading. We chose to fit exponentials to the H3K27me3 measurements to determine a simple statistic that reflects the degree of spreading. We then plotted the estimated decay lengths of the exponentials against the time points of the measurements (Fig. 2B & C). We find H3K27me3 spreads logarithmically over the six hours following mitosis 13, approaching an equilibrium (Fig. 2C). The derivative of the logarithmic fit to the decay lengths indicates H3K27me3 spreads at a rate of

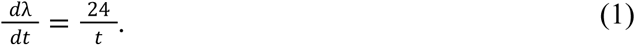

**Figure 2.**
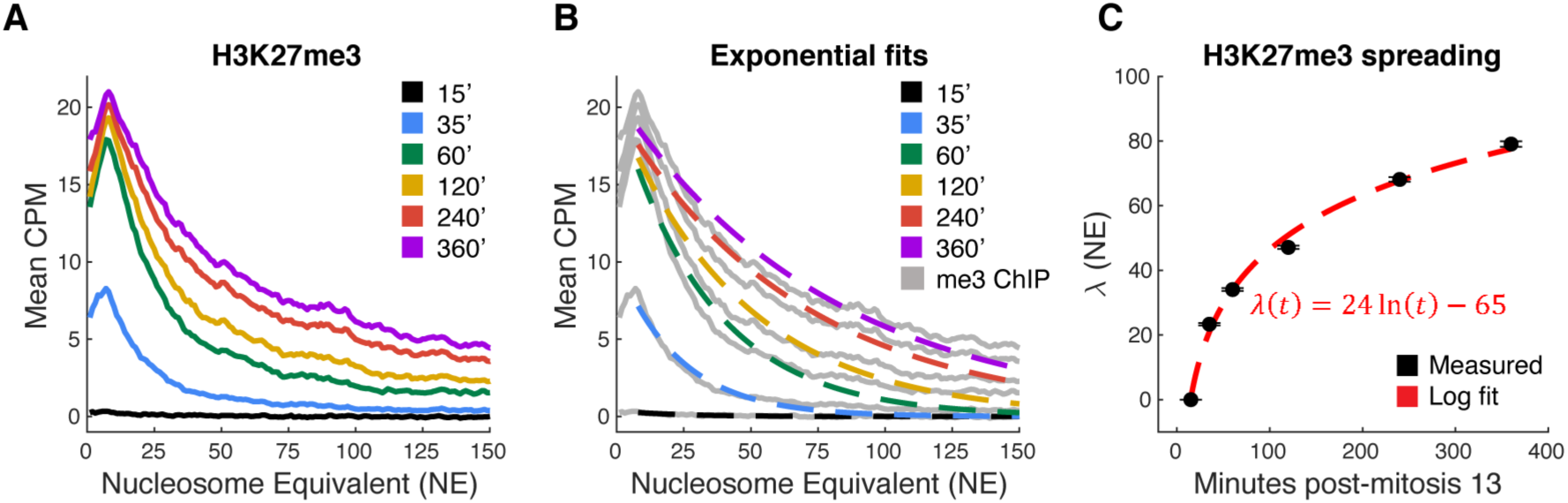
Quantification of *in vivo* H3K27me3 spreading. **A)** Mean CPM-normalized ChIP-seq measurements of H3K27me3 across isolated PRE halves, calculated while omitting measurements at nucleosomes past the first proximal boundary element to the E(z) peak (NE = 1). The legend indicates the minutes post-mitosis thirteen corresponding to each of the measurements. As in Fig. 1, NC14 + 15’, NC14 + 35’, and NC14 + 1h data is from an analysis of data collected in Gonzaga-Saavedra et al. (Gonzaga-Saavedra et al., 2026), while NC14 + 2h, NC14 + 4h, and NC14 + 6h data were collected in this work. **B)** Exponential fits to H3K27me3 measurements at isolated PREs over time. The legend indicates the minutes post-mitosis thirteen corresponding to the measurements that the exponentials were fit to. H3K27me3 averages are shown in grey. **C)** Decay lengths (λ) of the exponential curves presented in (B) plotted against the time points of the H3K27me3 measurements the exponentials were fit to. At NC14 + 15’, λ is set to 0, as at that time there is no measured H3K27me3 and therefore the exponential fit is uninterpretable. A log fit to these decay lengths indicates H3K27me3 spreads following mitosis 13 according to λ(*t*) = 24ln(*t*) - 65. Error bars indicate the standard errors of the decay length estimates.

Eq. 1 indicates that H3K27me3 spreads at an average rate of 0.43 nucleosomes/minute over the six hours following mitosis 13. This quantitative analysis can serve as a foundation for building a mathematical model to dissect the molecular mechanisms underlying the establishment of epigenetic silencing. Through its deposition of H3K27me3 on a chromatin template initially devoid of the modification, the early *Drosophila* embryo provides an *in vivo* context for determining reaction kinetics typically only quantified *in vitro*, with the additional benefit of spatially resolving how modifications spread over long genomic distances.

### A reaction-diffusion framework for Polycomb spreading

Modeling epigenetic systems requires making abstractions of often complex molecular interactions. PRC2 activity can be influenced by a variety of cofactors and DNA binding proteins (Brown et al., 1998, 2003, 2018; Horard et al., 2000; Klymenko et al., 2006; Ray et al., 2016; Tamburri et al., 2024; L. Wang et al., 2004), as well as the activity of Polycomb Repressive Complex 1 (PRC1) (Tamburri et al., 2024), which deposits ubiquitination at histone H2A lysine 118 (H2Aub), a mark that contributes to the recruitment and catalytic activity of PRC2 by interacting with the PRC2 accessory subunits Jarid2 (Cooper et al., 2016; Kasinath et al., 2021; Sanulli et al., 2015) and jing/AEBP2 (Kalb et al., 2014; Kasinath et al., 2021; Kim et al., 2009). Furthermore, the PRC2 subunit Extra Sex Combs (Esc, flies) / Embryonic Ectoderm Development (EED, mammals) binds to existing H3K27me2/3 to allosterically enhance the catalytic activity of E(z)/EZH2 (Margueron et al., 2009). The net effect of these molecular interactions is that, to establish Polycomb domains *de novo*, PRC2 is recruited to chromatin and then the activity of the complex progresses out from nucleation sites through a spreading mechanism that depends on Esc/EED (Højfeldt et al., 2018; Oksuz et al., 2018). Work has proposed that PRC2 can also interact with the genome through a non-specific “hit-and-run” mechanism that leads to accumulation of low levels of H3K27 methylation distal to nucleation sites (Oksuz et al., 2018). We sought to construct a mathematical model that quantitatively predicts H3K27me3 spreading along chromatin.

In defining the model, we refer to E(z) rather than PRC2, as we are interested in the catalytic behavior of the complex. We assume that the other PRC2 subunits are not concentration-limiting such that when E(z) associates with the genome it can readily methylate histones (Bonnet et al., 2019). We hypothesize that, due to both diffusion of E(z) and sub-diffusive fluctuations in chromatin fibers, the activity of E(z) progresses out from a nucleation site effectively according to a 1D random walk; E(z) likely methylates nucleosomes directly adjacent to a nucleation site sooner than it methylates more distal nucleosomes. As E(z) progresses along the genome (“diffusing”), it catalyzes a sequence of methylation reactions, upgrading H3K27me0 to-me1 to-me2 to-me3. Therefore, we chose to develop a reaction-diffusion framework for modeling the *de novo* establishment of Polycomb modifications surrounding a nucleation site.

We formulated a reaction-diffusion system with five state variables: E(z) and H3K27me0,-me1,-me2, and-me3 (Fig. 3A). In the framework, E(z) progresses along a chromatin fiber in 1D according to its diffusion coefficient *D*, while H3K27 methylation states lack the ability to diffuse and occupy fixed nucleosome coordinates (Fig. 3B). At every nucleosome position in a modeled array, the forward transitions [me0 → me1], [me1 → me2], and [me2 → me3] can occur, which all depend on the concentration of E(z) engaged with the substrate, and reflect catalysis of H3K27 methylation (Fig. 3A & B). These transition rates also depend on the parameter *R*, which represents the baseline catalytic activity of E(z) (nM^-1^ min^-1^), and is scaled according to the methylation substrate (Fig. 3A & B). *In vitro* measurements of EZH2 activity have indicated [me0 → me1]:[me1 → me2]:[me2 → me3] ratios of catalytic efficiencies are 13:4:1, as E(z)/EZH2 catalyzes lower-order methylation groups more quickly than higher-order groups (Sneeringer et al., 2010). We therefore chose to scale the forward methylation transitions in our model according to 13:4:1. Our model also incorporates reverse transitions between methylation states, reflecting mechanisms for active demethylation and histone turnover (Fig. 3A & B). H3K27me1,-me2, and-me3 groups can all transition to-me0 due to baseline histone turnover and the incorporation of unmodified histones to chromatin at DNA replication, with these transitions in the model dependent on the rate constant *d_H_* (min^-1^). Further, H3K27me2 and-me3 groups can be demethylated due to the activity of the H3K27 demethylase Utx (Smith et al., 2008). In the model, the reverse transitions due to active demethylation depend on the rate constant *d_UTX_*(min^-1^). While a parameter sweep revealed that the value of *d_UTX_*does not impact the model’s H3K27me3 predictions (Materials & Methods), consistent with experimental results that show a loss of Utx does not significantly affect H3K27me3 levels in *Drosophila* (Copur & Müller, 2013), we nonetheless incorporated the parameter into the model for completeness.

**Figure 3.**
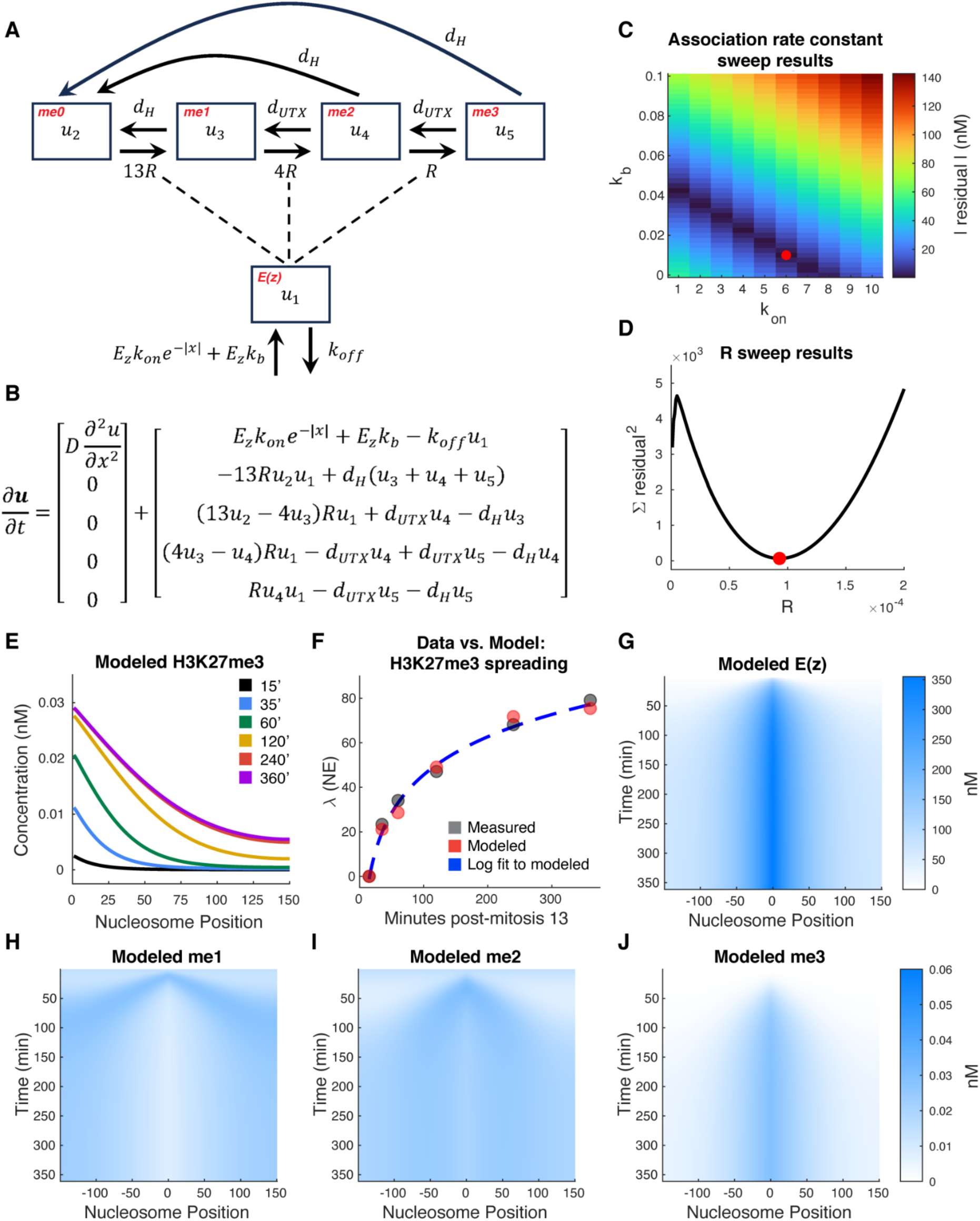
A reaction-diffusion system predicts *in vivo* H3K27me3 spreading. **A)** Compartment diagram of the reactions of the reaction-diffusion system. *u_1_* corresponds to the concentration of E(z) at a nucleosome position, and *u_2_*, *u_3_*, *u_4_*, and *u_5_* correspond to the concentrations of H3K27me0,-me1,-me2, and-me3 respectively at the nucleosome. The parameters governing each transition (*R*, *d_H_*, *d_UTX_*, *E_z_*, *k_on_*, *k_b_*, and *k_off_*) are described in the main text and Materials and Methods. **B)** PDEs that define the dynamics of the state variables in the reaction-diffusion model. E(z) (*u_1_*) “diffuses” along the nucleosome array according to the diffusion coefficient *D*. The rates of change of the H3K27 methylation states (*u_2_*, *u_3_*, *u_4_*, and *u_5_*) at each nucleosome are defined according to mass action kinetics. **C)** Parameter sweep results for the model’s E(z) nucleation rate constant, *k_on_*, and background E(z) association rate constant, *k_b_*. The heatmap shows, at each parameter combination, the absolute value of the difference between the predicted E(z) concentration across the nucleosome array over NC14 (t = 0 to 60 minutes) and 60 nM, the concentration of E(z) present in the nucleus calculated from experimental measurements (Bonnet et al., 2019; Degen et al., 2026; Gonzaga-Saavedra et al., 2026). x- and y-axis units are min^-1^. The best-fit parameter combination (*k_on_* = 6 min^-1^, *k_b_*= 0.01 min^-1^) is indicated by the red point. **D)** Parameter sweep results for *R*, the model’s baseline E(z) catalytic activity parameter. Plotted is the sum of squared residuals determined by comparing the decay lengths of the exponential fits to the model’s H3K27me3 predictions at *t* = 35, 60, 120, 240, and 360 minutes to the decay lengths of the exponential fits to the corresponding mean H3K27me3 measurements at isolated PREs shown in Fig. 2B. y-axis units are nucleosomes^2^, and x-axis units are nM^-1^ min^-1^. The best-fit *R* value (*R* = 9.3 x 10^-5^ nM^-1^ min^-1^) is indicated by the red point. **E)** Model predictions of H3K27me3 concentrations along the nucleosome array at *t* = 15, 35, 60, 120, 240, and 360 minutes. Nucleosome Position = 0 marks the site of E(z) nucleation. The model simulates the establishment of an H3K27me3 domain. **F)** Comparison of the decay lengths of exponential fits to the modeled H3K27me3 vs. measured H3K27me3 at isolated PREs (Fig. 2B & C). Decay lengths (λ) are plotted against the time points of the simulations/measurements. Measured and modeled λ’s at NC14 + 15’ are set to zero as *in vivo* H3K27me3 is absent from chromatin at this time. A log fit to the modeled λ (dashed blue line) indicates the model predicts the logarithmic spreading dynamics observed in the *in vivo* data. **G)** Modeled E(z) concentration along the 300-nucleosome array over 360 minutes. Nucleosome Position = 0 marks the site of E(z) nucleation. E(z) spreads out from the nucleation site and then reaches a steady-state where its concentration is enriched at the PRE and decays to a low baseline at farther nucleosomes. **H)** Modeled H3K27me1 concentration along the 300-nucleosome array over 360 minutes. Nucleosome Position = 0 marks the site of E(z) nucleation. The colorbar that applies to this panel is included in (J). The model simulates that, over time, H3K27me1 is depleted at the nucleation site and adjacent nucleosomes. **I)** Modeled H3K27me2 concentration along the 300-nucleosome array over 360 minutes. Nucleosome Position = 0 marks the site of E(z) nucleation. The colorbar that applies to this panel is included in (J). The model simulates that, over time, a broad domain of H3K27me2 forms. **J)** Modeled H3K27me3 concentration along the 300-nucleosome array over 360 minutes. Nucleosome Position = 0 marks the site of E(z) nucleation. The model simulates that, over time, a peaked domain of H3K27me3 forms centered on the E(z) nucleation site.

The model accounts for two mechanisms of E(z) association with the genome: nucleation at a PRE, and non-specific, hit-and-run activity. The rate of E(z) association with a nucleosome at position *x* in the modeled chromatin fiber is defined by

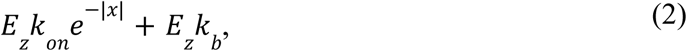

with *E_z_* representing the concentration of free E(z), *k_on_* the rate constant for E(z) nucleation on chromatin, and *k_b_* the rate constant for nonspecific E(z) association with the genome (Fig. 3A & B). At the PRE (*x* = 0), the rate of E(z) association is high (*E_z_k_on_*+*E_z_k_b_*). At nucleosomes farther from the PRE, the contribution of E(z) nucleation to the rate of E(z) association decreases exponentially according to *e^-\x\^*. Far from the PRE, the rate that E(z) associates with the genome is primarily determined by *E_z_k_b_*. Therefore, the model accounts for both nucleation-dependent E(z) recruitment to chromatin (which is high at PREs), as well as nucleation-independent E(z) association that is consistent with a hit-and-run mechanism. The model also allows for E(z) dissociation from the genome at any position *x* along the chromatin fiber dependent on the rate constant *k_off_* (min^-1^), the inverse of the average duration E(z) spends in the random walk (Fig. 3A & B).

**Figure Supplement 1.**
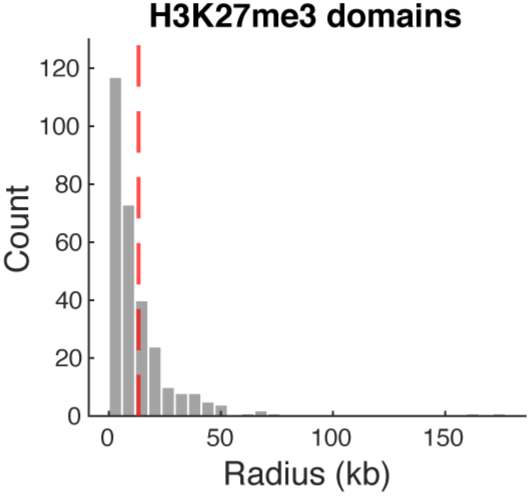
H3K27me3 domain radii. Plotted are the radii of H3K27me3 domains at NC14 determined previously from H3K27me3 ChIP-seq measurements (Gonzaga-Saavedra et al., 2026). The average domain radius of 13.3 kb is indicated by the dashed red line.

**Figure Supplement 2.**
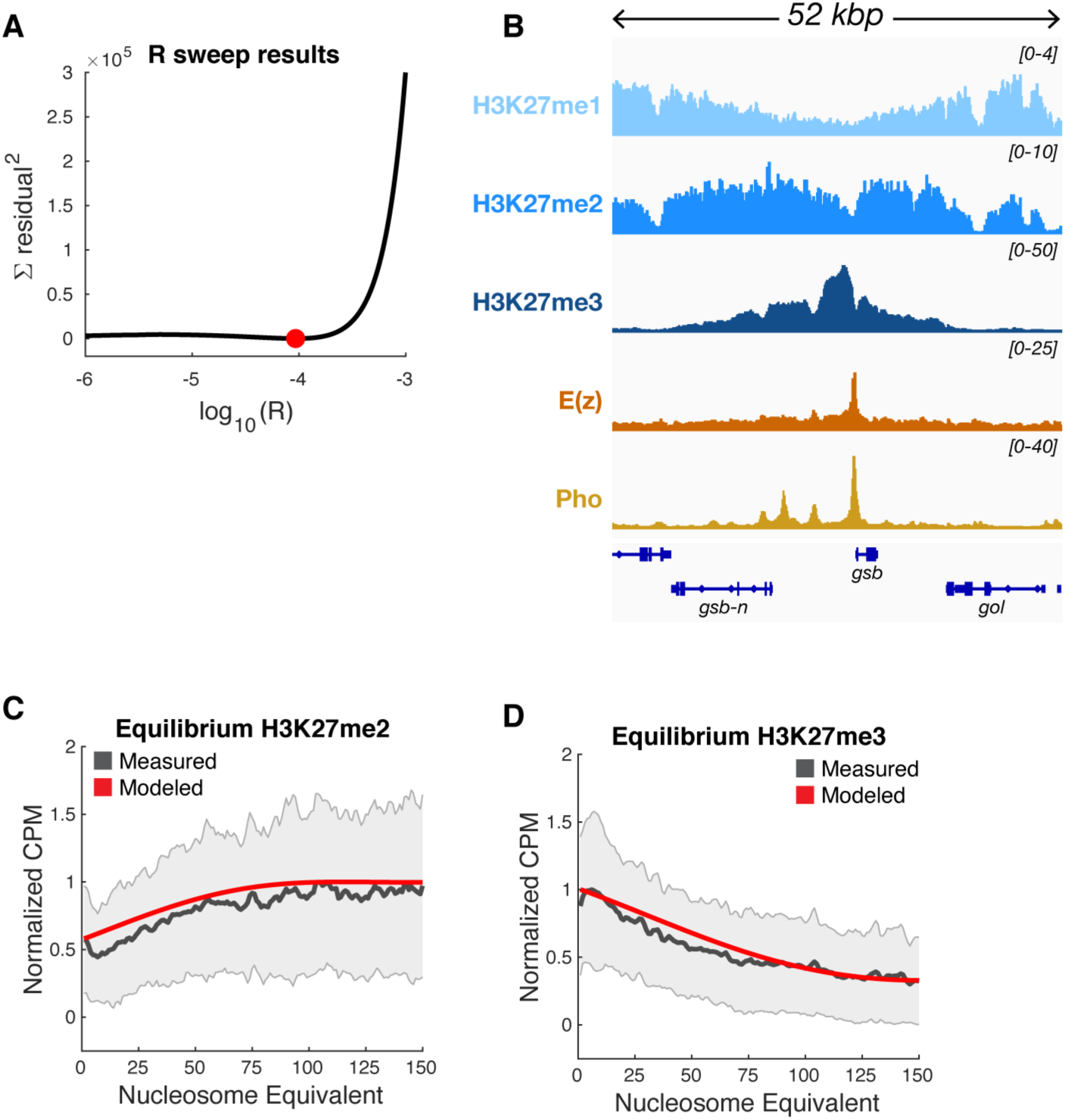
Parameter sweep results and equilibrium estimates. **A)** Full *R* parameter sweep results. As in Fig. 3D, plotted is the sum of squared residuals determined by comparing the decay lengths of the exponential fits to the model’s H3K27me3 predictions at *t* = 35, 60, 120, 240, and 360 minutes to the decay lengths of the exponential fits to the corresponding mean H3K27me3 measurements at isolated PREs shown in Fig. 2B. y-axis units are nucleosomes^2^, and x-axis units are log_10_(nM^-1^ min^-1^). The best-fit *R* value (*R* = 9.3 x 10^-5^ nM^-1^ min^-1^) is indicated by the red point. **B)** Experimental measurements of Polycomb modifications at a PRE at NC14 + 60’. Shown are ChIP-seq measurements of H3K27me1,-me2, and-me3 at the *gooseberry* (*gsb*) locus which contains a PRE occupied by E(z) and the nucleating factor Pleiohomeotic (Pho) at NC14. H3K27me1/2/3 were measured at NC14+60’, while E(z) and Pho occupancies were measured in broader NC14 embryo collections. H3K27me1, H3K27me3, E(z), and Pho data is from Gonzaga-Saavedra et al. (Gonzaga-Saavedra et al., 2026), and the H3K27me2 data is from Degen et al. (Degen et al., 2026). The depletion of me1, broad domain of me2, and peaked domain of me3 are consistent with the reaction-diffusion model’s predictions. **C)** Comparison of modeled vs. measured H3K27me2 at equilibrium. Plotted are mean ChIP-seq measurements of H3K27me2 calculated at isolated PREs while accounting for boundary elements (gray **±** standard deviation) (Materials and Methods). Experimental data is from ChIP-seq measurements of H3K27me2 in 24-hour old embryos from Bonnet et al (Bonnet et al., 2022). The model’s prediction at *t* = 22 hours (corresponding to 24-hour old embryos, as *t* = 0 in the model marks ∼2 hours into development) is shown in red. To allow for comparison of the shapes of the distributions, both modeled and measured data is normalized to their respective maximums across the 150 nucleosomes. The model simulates the depletion of me2 at the PRE (nucleosome equivalent = 0) seen *in vivo*. **D)** Comparison of modeled vs. measured H3K27me3 at equilibrium. Plotted are mean ChIP-seq measurements of H3K27me3 calculated at isolated PREs while accounting for boundary elements (gray **±** standard deviation) (Materials and Methods). Experimental data is from ChIP-seq measurements of H3K27me3 in 24-hour old embryos from Bonnet et al (Bonnet et al., 2022). The model’s prediction at *t* = 22 hours is shown in red. Both modeled and measured data is normalized to their respective maximums across the 150 nucleosomes. The model simulates the decay of H3K27me3 signal at nucleosomes farther from the PRE.

### The model predicts *in vivo* Polydomb dynamics

After defining the model, we sought to evaluate if it could recapitulate our *in vivo* measurements of H3K27me3 spreading along chromatin. The model contains eight parameters, five of which we fixed based on the literature or our measurements (Materials and Methods). A key parameter of the model is the diffusion coefficient *D*, that determines how quickly E(z) progresses along the chromatin fiber. As we didn’t explicitly account for the activity of the Esc/EED subunit of PRC2—known to facilitate PRC2 spreading from nucleation sites—the model in part captures this activity through its definition of *D*, which we calculated from the mean square displacement of the 1D random walk,

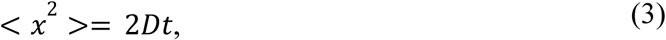

where *x* represents the displacement of PRC2 after diffusing in 1D with a diffusion coefficient *D* over a period of time *t*. We estimated the displacement of E(z) from nucleation sites during NC14 by drawing upon measurements of H3K27me3 domain widths after an average of 50 minutes post-mitosis 13 (Gonzaga-Saavedra et al., 2026). Given the average H3K27me3 domain radius of 13.3 kilobases from a central PRE (Fig. S1) and a nucleosome footprint of 180 basepairs, <*x*^2^> = 5,460 nucleosomes^2^, and thus

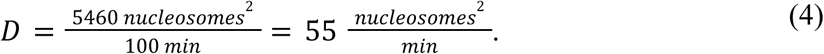

This diffusion coefficient equates to ∼3 x 10^4^ bp^2^ s^-1^ and is one to two orders of magnitude lower than 1D diffusion coefficients describing the sliding of transcription factors along DNA (Blainey et al., 2006; Davidson et al., 2023; Komazin-Meredith et al., 2008; Sakong et al., 2026; Tafvizi et al., 2008). The relatively slow rate of potential PRC2 progression along the genome may stem from nucleosomes acting as “sticky” substrates that interact highly with the complex. We use *D* = 55 nucleosomes^2^ min^-1^ for the wild-type model presented in the main text. However, as this parameter value approximates the result of likely a multitude of molecular events, we assess the model’s dependency on *D* in the Materials and Methods.

After defining parameters according to experimental measurements, the model contains three free parameters: the baseline E(z) catalytic activity (*R*), the rate constant for E(z) nucleation at a PRE (*k_on_*), and the rate constant for background E(z) association with the genome (*k_b_*). As *k_on_* and *k_b_* determine the concentration of E(z) that accumulates on the chromatin fiber, we calibrated these parameters such that the model predicts an average [E(z)] along the fiber during NC14 of ∼60 nM, the estimated [E(z)] in the nucleus at this time (Fig. 3C) (Degen et al., 2026). Over the tested parameters values, *k_on_* = 6 min^-1^ and *k_b_*= 0.01 min^-1^ best predict the nuclear E(z) concentration (Fig. 3C). *k_on_*= 6 min^-1^ closely approximates the effective rate constant for general E(z) association with chromatin measured via FRAP, *k*_on_* = 8 min^-1^ (Steffen et al., 2013). As expected, *k_b_* << *k_on_*, as the probability of E(z) recruitment to a nucleosome is much less when the nucleosome is far from a PRE versus at the PRE. To determine the best-fit value of *R*, we swept across *R* values, finding that the model best predicts the decay lengths of the measured H3K27me3 averages across timepoints at *R* = 9.3 x 10^-5^ nM^-1^ min^-1^ (Fig. 3D, Fig. S2 A).

With these parameter values, the reaction-diffusion framework simulates the establishment of a domain of H3K27me3 over six hours post-mitosis 13 (Fig. 3E). The model recapitulates the decay lengths of measured H3K27me3 at each time point, and thus the logarithmic spreading of the modification from PREs (Fig. 3F). From the logarithmic fit to the decay lengths of the simulated H3K72me3, the modeled H3K27me3 spreading rate is

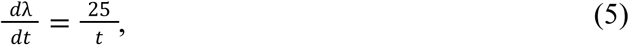

closely matching the spreading rate determined from the experimental data (Eq. 1). The success of the model supports our conceptualization of how E(z) effectively progresses along chromatin according to a 1D random walk.

In addition to simulating the canonical mark of gene silencing, H3K27me3, the model generates predictions for the spatiotemporal dynamics of E(z) and lower-order H3K27 methylation states on a chromatin fiber (Fig. 3G-J, Supplemental Video 1). The model predicts that E(z) accumulates at the PRE (*x* = 0) very quickly following the start of NC14 (*t* = 0), and then diffuses out bidirectionally along chromatin (Fig. 3G). At 30 minutes following mitosis 13, simulated H3K27me1 levels have dropped at the PRE, while a substantial amount of H3K27me2 has accumulated (Fig. 3H & I, Supplemental Video 1). By two hours post-mitosis 13, the model anticipates a lack of H3K27me1 close to the PRE and a broad distribution of H3K27me2 (Fig. 3H & I, Supplemental Video 1). By this point, a peaked domain of H3K27me3 is also simulated (Fig. 3J, Supplemental Video 1). The equilibrium that the simulation settles into—showing a depletion of me1 close to the PRE, a broad distribution of me2, and a more focal domain of me3—is consistent with trends seen in experimental data (Fig. 3G-J, Fig. S2 B, Supplemental Video 1) (Bonnet et al., 2022; Gonzaga-Saavedra et al., 2026). Over the course of NC14, H3K27me1 decreases at PREs (Gonzaga-Saavedra et al., 2026). Further, in late-stage *Drosophila* embryos, H3K27me2 covers chromatin in broad domains, while H3K27me3 is distributed in peaks at PREs (Bonnet et al., 2022).

We next evaluated how well the model predicts the equilibrium distributions of H3K27me2 and H3K27me3 by drawing upon ChIP-seq measurements of these modifications in 24 hour-old *Drosophila* embryos (Bonnet et al., 2022). At this late stage of embryogenesis, Polycomb domains have completed their establishment and have entered a phase of stable maintenance. To allow for comparison to model outputs, we analyzed the 24h H3K27me2 and-me3 measurements at isolated PREs delimited by domain boundaries. In 24h embryos, at isolated PREs, H3K27me3 is enriched, while H3K27me2 is depleted (Fig. S2 C & D). Farther from the PRE, H3K27me3 levels decrease, while H3K27me2 levels increase (Fig. S2 C & D). The equilibrium predicted by the model reflects these patterns (Fig. S2 C & D) (Materials and Methods). The ability of the model to predict steady-state patterns of H3K27me2/3 provides further support for the model’s assumptions. While the Polycomb system involves interactions between a dozen or more conserved PcG proteins, a minimal system where E(z) associates with chromatin primarily at nucleation sites and then effectively performs a 1D random walk to catalyze a sequence of methylation reactions is sufficient to recapitulate experimental observations.

### PRC1 activity modulates E(z) reaction rates and genomic association

Our model does not explicitly account for the role of PRC1 in regulating the activity of PRC2/E(z) by depositing H2Aub. H2Aub interacts with Jarid2, an accessory subunit of PRC2 that can recruit PRC2 to PRC1-modified sites, and when methylated, bind to Esc/EED to to lead to allosteric enhancement E(z)/EZH2 activity (Cooper et al., 2016; Kalb et al., 2014; Sanulli et al., 2015). Jarid2 can also negatively influence the catalytic activity of E(z)/EZH2 (Peng et al., 2009). In addition to Jarid2, H2Aub interacts with Jing/AEBP2, an accessory subunit of PRC2 that targets the complex to chromatin (Kalb et al., 2014; Kasinath et al., 2021; Kim et al., 2009). We hypothesized that H2Aub influences the parameterization of E(z) activity such that it speeds the rate H3K27me3 spreads along the genome. To determine how H2Aub quantitatively contributes to Polycomb spreading, we drew upon published ChIP-seq measurements (Bonnet et al., 2022) of H3K27me3 in 0-6h old wild-type *Drosophila* embryos and embryos mutant for *sex combs extra* (*Sce*), the catalytic subunit of PRC1 that deposits H2Aub (H. Wang et al., 2004). In embryos with the Sce[I48A] mutation that ablates Sce catalytic activity, lowered H3K27me3 levels accompany the depletion of H2Aub (Bonnet et al., 2022).

We analyzed the wild-type and Sce[I48A] datasets using our isolated PRE approach, which revealed loss of H2Aub leads to a substantial drop in H3K27me3 at isolated PREs (Fig. 4A). By fitting exponentials to the Sce[I48A] and control H3K27me3 averages, we quantified how the reduction in Sce activity decreases both the amplitude and spread of the H3K27me3 distribution at PREs (Fig. 4A). Compared to the wild-type distribution, the amplitude of H3K27me3 drops by 39% and the spread by 63% in Sce[I48A] mutants. We sought to determine which model parameters are most affected by loss of Sce catalytic function.

**Figure 4.**
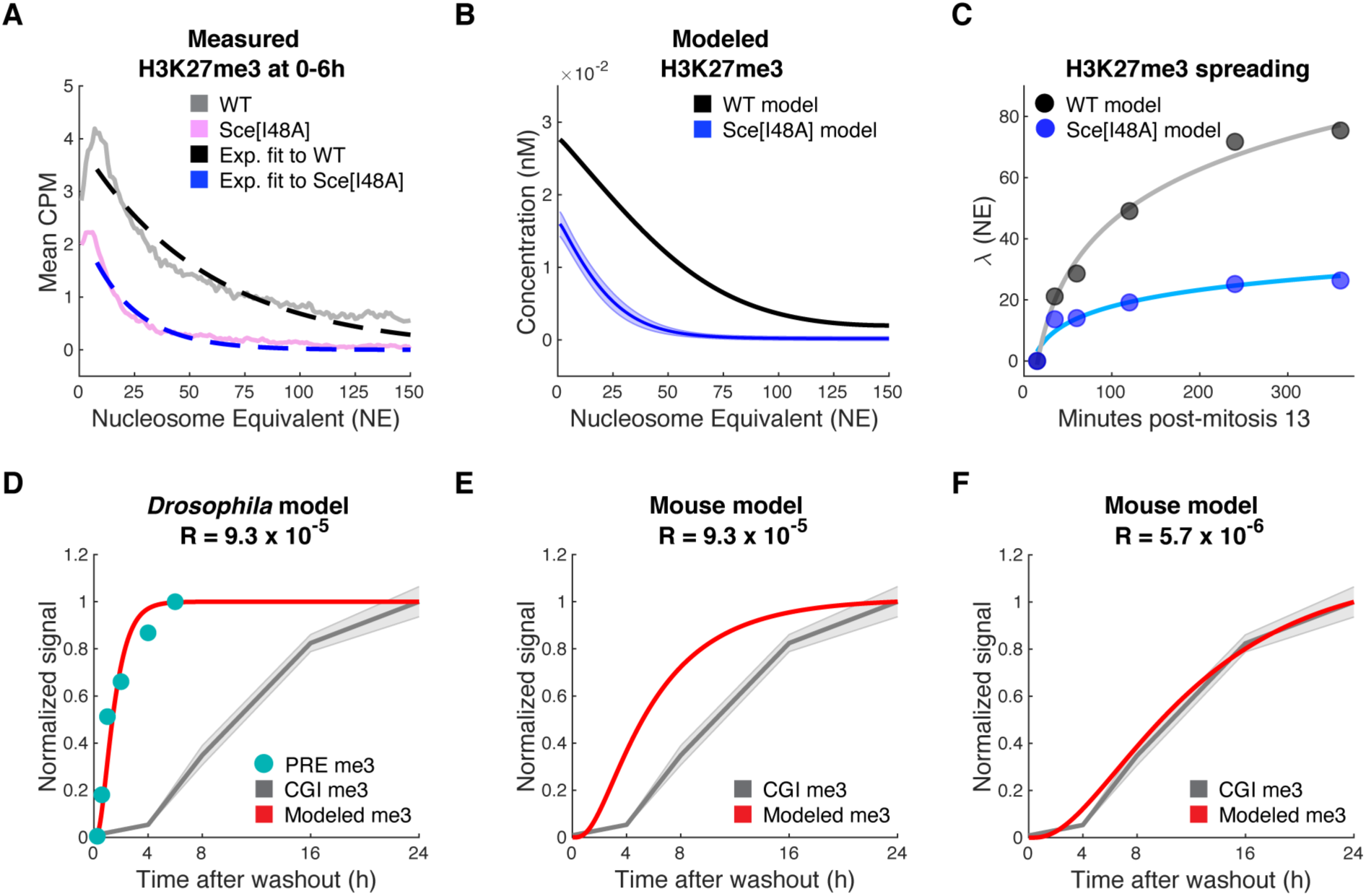
Simulating H3K27me3 in varied biological conditions. **A)** Diminished H3K27me3 at isolated PREs in Sce[I48A] mutants. Shown are mean CPM-normalized ChIP-seq measurements of H3K27me3 in wild-type (WT, gray) and Sce[I48A] (pink) embryos across isolated PREs, accounting for higher-order chromatin structure as defined in Materials and Methods. We performed our analysis on WT and Sce[I48A] measurements in 0-6h old embryos (timing from the onset of development) from Bonnet et al. (Bonnet et al., 2022). The exponential fit to the WT data is shown in black, and the exponential fit to the Sce[I48A] data is shown in blue. **B)** Model predictions of H3K27me3 at NC14 + 120’ with WT (black) and Sce[I48A] mutant (blue) parameter sets. As ChIP-seq measurements of H3K27me3 in 0-6h old embryos will enrich for sequencing reads from NC14 + 0’ to NC14 + 240’, the post-cleavage timeframe covered in the collection, we chose to model H3K27me3 at the average of this timeframe, NC14 + 120’. The “WT model” corresponds to the reaction-diffusion system when *k_on_* = 6 min^-1^, *k_b_* = 0.01 min^-1^, *R* = 9.3 x 10^-5^ nM^-1^ min^-1^, and *D* = 55 nucleosomes^2^ min^-1^, and the “Sce[I48A] model” corresponds to the reaction-diffusion system with *k_on_*, *k_b_*, *R*, and *D* values defined according to the top 1% parameters sets that allow for prediction of the mutant measurements. All other parameters are set at their baseline values defined in Materials and Methods. **C)** Modeled H3K27me3 spreading in wild-type and Sce[I48A] embryos. Decay lengths (λ, black and blue points) were determined by fitting exponentials to the model’s H3K27me3 predictions at NC14 +35’, +60’, +120’, +240’, and +360’. The “WT model” and “Sce[I48A] model” are as described in (B). At NC14 + 15’, the decay lengths of the WT and Sce[I48A] models were set to 0, as at that time H3K27me3 is not detectable experimentally and therefore the measured λ is uninterpretable (Fig. 2B). Logarithmic fits to the modeled λ are shown in gray (WT) and light blue (Sce[I48A]). The reaction-diffusion framework predicts that H3K27me3 spreads more slowly in the absence of H2Aub. **D)** Comparison of the H3K27me3 predictions from the *Drosophila*-configured reaction-diffusion model with H3K27me3 measurements at CpG Islands (CGIs) from Højfeldt et al (Højfeldt et al., 2018). Shown in red is the normalized integrated H3K27me3 concentration that the model predicts along a 300 nucleosome array over 24 hours post-mitosis 13 (*t* = 0). Cyan points indicate the normalized integrated H3K27me3 signal at isolated PREs calculated from the ChIP-seq measurements at NC14 + 15’, NC14 + 35’, and NC14 + 60’ made in Gonzaga-Saavedra et al. (Gonzaga-Saavedra et al., 2026), and the measurements at NC14 + 120’, NC14 + 240’, and NC14 + 360’ from this work. Shown in gray (± standard deviation) is the averaged normalized ChIP-qPCR signal across CGIs post-EZH2 inhibitor washout calculated from measurements made in Højfeldt et al (Højfeldt et al., 2018). The *Drosophila*-configured model recapitulates the dynamics of H3K27me3 establishment at PREs, but not the dynamics of establishment CGIs. **E)** Comparison of the H3K27me3 predictions from the mouse-configured reaction-diffusion model (in this case using *R* = 9.3 x 10^-5^ nM^-1^ min^-1^) with H3K27me3 measurements at CGIs from Højfeldt et al (Højfeldt et al., 2018). Shown in red is the normalized integrated H3K27me3 concentration that the “mouse” model (with *R* = 9.3 x 10^-5^ nM^-1^ min^-1^) predicts along a 300 nucleosome array over 24 hours post-EZH2 inhibitor washout (*t* = 0). As in (D), shown in gray (± standard deviation) is the averaged normalized ChIP-qPCR signal across CGIs calculated from measurements made in Højfeldt et al (Højfeldt et al., 2018). With *R* = 9.3 x 10^-5^ nM^-1^ min^-1^, the mouse model fails to recapitulate the slow accumulation of H3K27me3 post-EZH2 inhibitor washout. **F)** Comparison of the H3K27me3 predictions from the mouse-configured reaction-diffusion model (in this case using *R* = 5.7 x 10^-6^ nM^-1^ min^-1^) with H3K27me3 measurements at CGIs from Højfeldt et al (Højfeldt et al., 2018). Shown in red is the normalized integrated H3K27me3 concentration that the “mouse” model (with *R* = 5.7 x 10^-6^ nM^-1^ min^-1^) predicts along a 300 nucleosome array over 24 hours post-EZH2 inhibitor washout (*t* = 0). As in (D) and (E), shown in gray (± standard deviation) is the averaged normalized ChIP-qPCR signal across CGIs calculated from measurements made in Højfeldt et al (Højfeldt et al., 2018). With *R* = 5.7 x 10^-6^ nM^-1^ min^-1^, the mouse model recapitulates the slow accumulation of H3K27me3 post-EZH2 inhibitor washout.

We applied the reaction-diffusion framework to simulate the effects of mutating Sce on H3K27me3 patterns by sweeping across values of *k_on_*, *k_b_*, *R*, and *D*, the model’s parameters that describe E(z) recruitment, catalytic activity, and progression along the genome. With each parameter combination, we evaluated how well the model predicted the relative changes in the amplitude and spread of H3K27me3 produced by the Sce[I48A] mutation (Materials and Methods). We found that the model accurately predicts the H3K27me3 distribution in Sce[I48A] embryos by decreasing *k_on_*, *k_b_*, *R*, and *D* (Fig. 4B, Fig. S3). In the top 1% of parameter sets that best predict H3K27me3 in Sce[I48A], for each parameter, the majority of values fall below the wild-type value (Fig. S3). The results of this sweep are further detailed in Materials and Methods. With the mutant parameter sets, the model recapitulates the trends of the experimental measurements (Fig. 4A & B), simulating a 39% reduction in the amplitude and 61% reduction in the spread of H3K27me3, comparable to the 39% and 63% reductions determined by comparing the exponential fits of the Sce[I48A] and control H3K27me3 measurements.

These parameter sweep results are consistent with prior experimental findings that indicate the presence of H2Aub on chromatin increases the rates of E(z)/EZH2 recruitment and catalysis of H3K27 methylation (Blackledge et al., 2014, 2020; Kalb et al., 2014; Kasinath et al., 2021; Ohtomo et al., 2023; Tamburri et al., 2020), as the model predicts that E(z) has higher *k_on_*, *k_b_*, and *R* values in wild-type embryos than Sce[I48A] embryos. Notably, our results also indicate that H2Aub increases the speed at which E(z) samples histones along the chromatin fiber, which is represented by the 1D diffusion coefficient, *D*. The model predicts *D* is substantially higher in wild-type embryos than Sce[I48A] embryos. Further, compared with the broad distributions of the other parameters, *D* values are tightly distributed in the top 1% of mutant parameter sets, indicating that this parameter strongly influences prediction of the amplitude and spread of H3K27me3 in Sce[I48A] embryos (Fig. S3). Together, our parameter sweep results support prior work, while suggesting an additional and important role for H2Aub: to facilitate PRC2’s sampling of its histone substrates, independently of the complex’s catalytic rates and recruitment to chromatin. Future work will address how the PcG proteins that interact with H2Aub influence the progression of PRC2 activity along chromatin.

With the mutant parameter settings, the model predicts that H3K27me3 spreads more slowly in Sce[I48A] than in wild-type conditions (Fig. 4C). The derivative of the logarithmic fit to the modeled Sce[I48A] decay lengths indicates that, in the absence of H2Aub, H3K27me3 spreads at a rate of

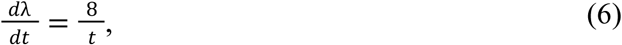

significantly slower than the spreading rate inferred from wild-type H3K27me3 measurements (Eq. 1). Thus, we propose that in wild-type conditions H2Aub contributes to E(z) activity such that it speeds the spread of H3K27me3 along chromatin.

**Figure Supplement 3.**
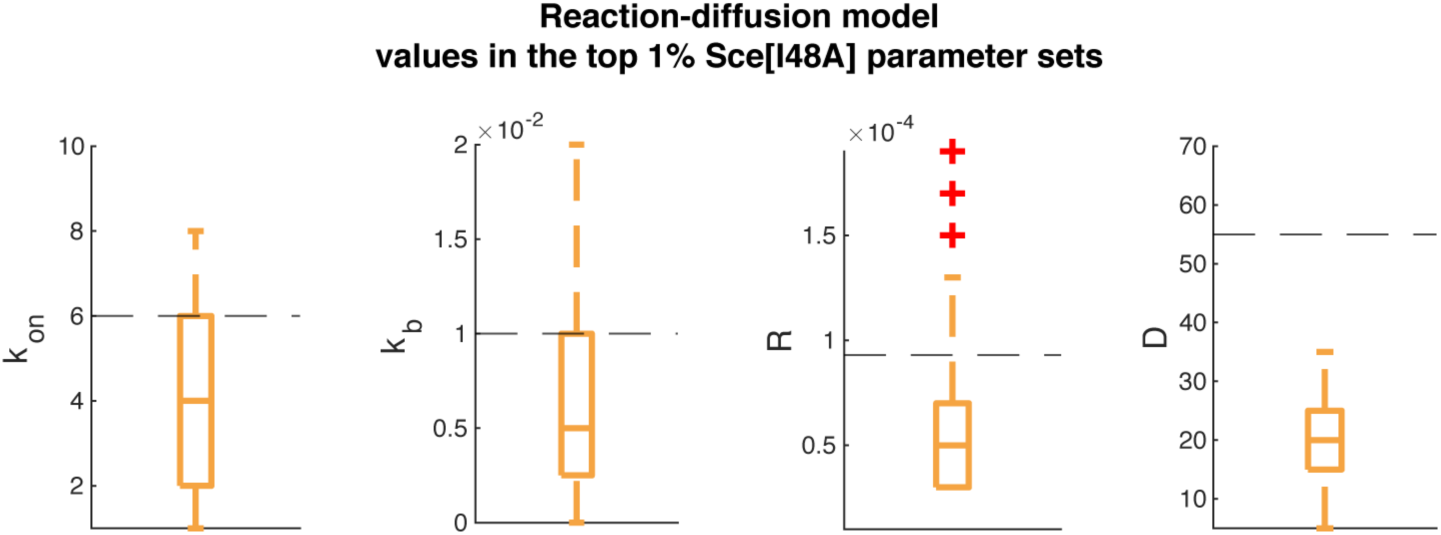
Parameters that best predict H3K27me3 in Sce[I48A] embryos. Shown are the distributions of values in the top 1% parameter sets (126 sets in total) that allow the reaction-diffusion model to best predict the decrease in magnitude and decay length of the exponential fits to the H3K27me3 measurements in WT and Sce[I48A] embryos. y-axes of the *k_on_* and *k_b_*plots have units of min^-1^, the y-axis of the *R* plot has units of nM^-1^ min^-1^, and the y-axis of the *D* plot has units of nucleosomes^2^ min^-1^. Dashed lines indicate the parameter values that allow the model to best predict the WT time course data as determined from parameter sweeps and experimental estimates (*k_on_* = 6 min^-1^, *k_b_* = 0.01 min^-1^, *R* = 9.3 x 10^-5^ nM^-1^ min^-1^, *D* = 55 nucleosomes^2^ min^-1^).

### The embryo as a super-charged environment for Polycomb activity

We next sought to investigate the significance of the baseline catalytic activity parameter in the model, *R*. In our construction of the model, we did not account for the specifics of how E(z) deposits methylation groups on histones. Instead, we defined *R* as a constant with units of nM^-1^ min^-1^ to allow for calculating the production rate of methylation groups given simple upgrading transitions and the law of mass action. Here, we provide a biochemical definition for this parameter, and evaluate its significance for interpreting how epigenetic states are regulated in the early embryo.

Assuming E(z) catalyzes H3K27me3 according to Michaelis-Menten reaction kinetics (Michaelis & Menten, 1913), the production rate of H3K27me3 at a given nucleosome can be calculated with

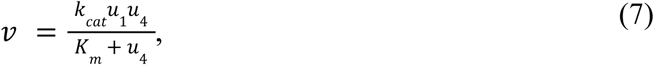

where *u_4_* represents the concentration of H3K27me2 at the nucleosome of interest, *u_1_* represents the total amount of E(z) engaged with the nucleosome, *k_cat_* represents the rate constant for H3K27me3 catalysis, and *K_m_* signifies the Michaelis constant of the reaction. Equating this expression to the the H3K27me3 production rate in our model, *Ru_4_u_1_*,

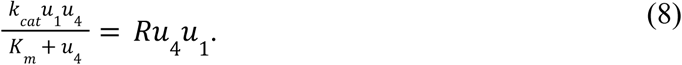

This equation can be simplified to calculate *R*:

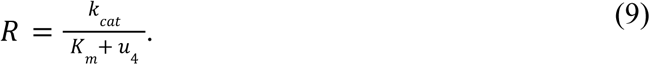

Given the concentration of any methylation group at a nucleosome is at most 0.06 nM (Materials and Methods) and *in vitro* experimental measurements indicate the Michaelis constants associated with E(z) catalytic activity are on the order of 100 nM (Sneeringer et al., 2010), *K_m_* >> *u_4_*, and therefore

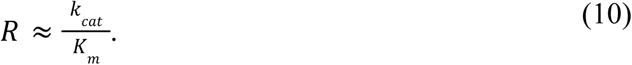

Thus, in the reaction-diffusion framework, *R* approximates the efficiency with which E(z) catalyzes the H3K27me2-me3 reaction. The basal efficiency by which E(z)/EZH2 catalyzes H3K27me3 has been measured *in vitro* to be 5.7 x 10^-6^ nM^-1^ min^-1^ (Sneeringer et al., 2010), while the value of *R* that best fits our experimental H3K27me3 time course data is *R* = 9.3 x 10^-5^ nM^-1^ min^-1^, 16 times higher than this basal efficiency. Our model therefore suggests that, in the early embryo, E(z) operates with elevated catalytic efficiencies.

We hypothesized that the basal catalytic efficiency of E(z) applies to cell culture and somatic contexts, rather than the early embryo. Therefore, we sought to evaluate if setting *R* to the basal catalytic efficiency would allow the model to predict the *de novo* establishment of H3K27me3 in mouse embryonic stem cells (mESCs), where H3K27me3 establishment has been measured previously (Højfeldt et al., 2018; Oksuz et al., 2018). By treating mESCs with an EZH2 inhibitor to deplete H3K27me2/3 and then washing away the inhibitor, a prior study tracked the re-establishment of H3K27 methylation on chromatin with ChIP (Højfeldt et al., 2018). We drew upon this study’s measurements of H3K27me3 dynamics at CpG islands (CGIs) in the mouse genome, DNA regions of high CG density at which PRC2 is nucleated. At CGIs, H3K27me3 increases on chromatin for 24 hours post-inhibitor washout (Højfeldt et al., 2018). We first tested whether the model, as configured for *Drosophila*, could predict these dynamics. As expected, with parameters tailored to *Drosophila* conditions, the reaction-diffusion framework fails to recapitulate the slow rise in H3K27me3 over 24 hours following inhibitor washout (Fig. 4D).

The *Drosophila* model predicts much more rapid establishment, consistent with our isolated PRE measurements (Fig. 4D). Therefore, we revised the model to tailor its parameters to mouse conditions.

To adjust the model to make mESC predictions, we defined the baseline demethylation rate constant *d_H_* given the cell cycle length and histone turnover rate in mESCs (Deaton et al., 2016; Waisman et al., 2019), and incorporated enzyme-mediated reverse transitions from me3 to me2, me2 to me1, and me1 to me0 according to effective rate constants determined in mammalian cells (Y. Zheng et al., 2012) (Materials and Methods). Further, we adjusted the E(z)/EZH2 concentration in the model to more accurately reflect the mammalian context (Y. Zheng et al., 2012). We left *k_off_*, *k_on_*, *k_back_*, and *D* at their *Drosophila* values, hypothesizing that E(z)/EZH2 similarly interacts with chromatin in regions of PRC2 nucleation in mESCs compared with PREs. With the “mouse” configuration of the reaction-diffusion framework and the best-fit *Drosophila* value of 9.3 x 10^-5^ nM^-1^ min^-1^, the model predicts a slower rate of H3K27me3 increase and delayed steady-state compared with the entirely *Drosophila*-tailored model (Fig. 4D & E). While the model adjustments allowed for closer approximation of the mESC *in vivo* H3K27me3 estimates, the model’s output still substantially over-shoots the rate of H3K27me3 re-establishment in mESCs (Fig. 4E).

We therefore lowered the E(z)/EZH2 baseline reaction efficiency in the mouse-tailored model to *R* = 5.7 x 10^-6^ nM^-1^ min, the measured basal efficiency with which EZH2 catalyzes H3K27me3 (Sneeringer et al., 2010). With the basal efficiency, the mouse model now approximates the rate of normalized H3K27me3 increase at CGIs (Fig. 4F). Linear fits to the measured and modeled H3K27me3 over the first 16 hours following *t* = 0 reveals the model’s predicted slope only deviates from the slope of the experimental measurements by 5.8%. The substantial improvement in the model’s prediction that resulted from lowering *R* to match the basal measurement suggests that, unlike the early *Drosophila* embryo, mammalian cell culture conditions do not significantly elevate E(z)/EZH2 catalytic activity. Additional H3K27me3 time course measurements in other embryonic contexts could test whether super-charged E(z)/EZH2 activity is a general feature of early embryogenesis.

## DISCUSSION

Here, we quantify the rate that H3K27me3 spreads *de novo* along the genome during *Drosophila* embryogenesis, and build a computational framework that predicts these dynamics. The model macroscopically represents the activity of a host of PcG proteins and cofactors, addressing the systems biology challenge of interpreting how a complex network of molecular interactions dictates silent chromatin states. We find that Polycomb dynamics in the early embryo are remarkably fast, and can be captured quantitatively by three key features of the system: the set of reactions that upgrade and downgrade methylation groups, the preferential association of E(z) with a nucleation site on chromatin, and the 1D diffusion of E(z) along the chromatin fiber. Through modeling the results of experimental perturbation to the Polycomb system, we discover that, in addition to modulating the catalytic rate and the genomic association of E(z) as expected, the Polycomb modification H2Aub likely facilitates the 1D random walk of E(z) along chromatin. We further dissect the Polycomb conditions of the early embryo by testing our model’s ability to predict H3K27me3 measurements in mammalian cell culture, and provide evidence that E(z) operates with an elevated catalytic efficiency in the earliest stages of embryogenesis. This work lays the groundwork for decoding how Polycomb spreading may dictate the dynamics of gene silencing across biological contexts.

Our work raises the hypothesis that the early embryonic chromatin environment enhances the catalytic activity of E(z)/EZH2 relative to more mature contexts. Across organisms, Polycomb modifications are passed down from the embryo’s parents, and could support epigenetic reprogramming in the zygote (Burton & Torres-Padilla, 2010; Kaneshiro et al., 2019; Zenk et al., 2017; H. Zheng et al., 2016). At later embryonic stages, the catalytic efficiency of E(z)/EZH2 may decline, as mature canonical H3K27me3 domains generally lack H3K27me2 and H2Aub (Bonnet et al., 2022; Ferrari et al., 2014; Højfeldt et al., 2018; Oksuz et al., 2018). Further, embryonic chromatin contains patterns of H3K36me3 and H3K4me3 (Li et al., 2014; Liu et al., 2016; Xu et al., 2019; Zhang et al., 2016; Zhu et al., 2019), modifications that antagonize the deposition of H3K27 methylation (Cookis et al., 2025; Schmitges et al., 2011; Y. Zheng et al., 2012). How the overall epigenetic landscape programs *in vivo* Polycomb dynamics throughout developmental stages remains unknown. In this work, we provide the necessary sampling of H3K27me3 and computational analysis to illustrate that the baseline catalytic efficiency for E(z) is enhanced in early embryonic contexts, providing a molecular mechanism for the prior qualitative observation that Polycomb modifications accumulate quickly in embryos. While H3K27me3 measurements in additional embryonic systems suggest this mechanism is generalizable across organisms (Akkers et al., 2009; Hickey et al., 2022; Matsuwaka et al., 2025; van Heeringen et al., 2014), sampling H3K27me3 at higher rates in these systems is required to make a definitive conclusion. In addition, it is of interest that the reaction-diffusion model predicts H3K27me3 re-establishment dynamics in mESCs using the basal catalytic efficiency of EZH2, implying that in mESCs, the enzyme does not rely on pre-existing histone modifications to boost its activity. How E(z)/EZH2 achieves a higher baseline catalytic efficiency in early embryos is currently unclear. While differences in histone modification states could underlie the differences in E(z)/EZH2 activity between embryos and cultured cell systems, additional factors, such as post-translational modification of the enzyme, could also contribute.

We provide support for H2Aub acting to facilitate the formation of H3K27me3 domains during early embryogenesis. The model predicts that, in addition to modulating the rates at which E(z) is nucleated on chromatin and catalyzes methylation, Sce/H2Aub speeds the random walk of PRC2 along the genome and thus the spreading of H3K27me3. The *Drosophila* embryo may rely on H2Aub to assist H3K27me3 establishment, as H2Aub pre-coats Polycomb domains at the start of NC14, with the mark largely overlapping the H3K27me3 domains that form over the nuclear cycle (Gonzaga-Saavedra et al., 2026). Whether the Polycomb spreading rate potentially programmed by Sce/H2Aub determines gene expression dynamics remains to be determined. RNA-seq and immunostaining experiments have indicated that the Sce[I48A] mutation does not significantly impact Homeotic (Hox) gene expression levels (Bonnet et al., 2022; Pengelly et al., 2015). It is possible that the presence of multiple PRC2 nucleation sites at the Hox genes mitigates the dependence of gene expression on HK27me3 spreading rate. However, measurements of gene expression dynamics at additional loci may reveal Polycomb spreading rates have phenotypic impacts. In support of a role for H2Aub—and possibly H3K27me3 spreading rate—in cell fate specification in the embryo, prior work has shown Jarid2 is required for the establishment of gene silencing during the differentiation of embryonic stem cells to neural progenitors (Petracovici & Bonasio, 2021). While Jarid2 likely plays varied roles in PRC2-mediated gene regulation in *Drosophila* (Herz et al., 2012), Jing/AEBP2 could also contribute to gene silencing by facilitating Polycomb spreading at regions marked with H2Aub. Further work is required to determine the relationship between Polycomb kinetics and gene expression.

The success of the reaction-diffusion framework in recapitulating *in vivo* Polycomb states supports a conceptualization of how, after recruitment to nucleation sites, PRC2/E(z) diffuses in 1D along chromatin. Fluorescence recovery after photobleaching measurements have indicated that in the early *Drosophila* embryo, E(z) has an average residence time on chromatin of 1.52 seconds (Steffen et al., 2013). The transient nature of E(z) contacts with the genome suggest PRC2 might not remain stably associated with PREs. Instead, nucleation sites may define the genomic locations where E(z) transitions from 3D to 1D diffusion. The possibility exists however, that E(z) has a longer residence time and is “tethered” at PREs, instead relying on chromatin fluctuations to contact nucleosomes along the fiber. Indeed, prior work modeling H3K27 methylation in the *Drosophila* embryo accounted for potential PRE tethering by approximating chromatin contact probabilities (Lundkvist et al., 2023). As the diffusion coefficient of our model is calculated from *in vivo* data and macroscopically represents a collection of molecular events, this parameter has the capacity to account for close-range chromatin contacts within TADs bringing nucleosomes in proximity to E(z).

Our analysis of H3K27me3 at isolated PREs in relationship to boundary elements highlights a link between PRC2 activity and higher-order chromatin structure. The correlation we see between TAD boundary elements and H3K27me3 domain boundaries suggests that higher-order chromatin organization limits the extent of Polycomb spreading. Prior work has also indicated that PcG proteins facilitate H3K27 methylation spreading by regulating chromatin contacts (Kraft et al., 2022; Ogiyama et al., 2018; Sabarís et al., 2026). PRC2 might rely on long-range 3D contacts to spread H3K27me2/3 in mESCs (Oksuz et al., 2018), and computational modeling has shown 3D genome folding likely contributes to the epigenetic memory of heterochromatin domains (Owen et al., 2023). Future studies will address the feedback mechanisms between the Polycomb system and genome architecture.

Going forward, the model can be used as a tool to quantitatively define the impact of both PcG proteins and higher-order chromatin structure on the establishment of Polycomb domains. With the development of approaches to perturb chromatin organization, the model can help us decode whether chromatin contacts contribute to the 1D progression of E(z) activity within TADs. Furthermore, with additional H3K27me3 spreading measurements in early developmental systems, which can be made through standard genome sequencing approaches, the model can be fit to explore the commonalities and distinctions in Polycomb conditions across organisms.

## MATERIALS AND METHODS

### EXPERIMENTAL MODEL DETAILS

#### Drosophila stock

*y w; RpA70-EGFP* flies (Blythe & Wieschaus, 2015) produced the embryos used for our ChIP-seq measurements of H3K27me3.

### METHOD DETAILS

#### Chromatin immunoprecipitation

##### Fixation

As described previously (Degen et al., 2026), embryos were dechorionated in 4% Sodium Hypochlorite (Clorox) and washed extensively in deionized water. Following dechorionation, embryos were fixed for 15 minutes in 6 ml Heptane, 2 ml PBS + 0.5% Triton X-100, and 180 µl 20% Formaldehyde. Fixation was quenched by incubating the embryos in PBS + 0.5% Triton X-100 + 125 mM Glycine for two minutes. Embryos were then washed thrice in ice-cold PBS + 0.5% Triton X-100.

##### Embryo sorting

Fixed embryos were sorted under PBS + 0.5% Triton X-100 in 35 mm petri dishes lined with 1% agarose. Embryos were collected at stage 9 (∼NC14 + 2h), stage 11 (∼NC14 + 4h), and stage 12 (∼NC14 + 6h). Stage 9 was defined as “just finished germband extension, no visible stomodeal invagination,” stage 11 was “not yet retracting germband, clear stomodeal invagination, visible parasegments,” and stage 12 was “beginning/early germband retraction.” For each stage, embryos were collected for two ChIP-seq replicates (20 embryos per replicate). Sorted embryos were then frozen at-80°C until thawing immediately prior to the ChIP-seq protocol.

##### Chromatin Immunoprecipitation

ChIP-seq was performed as described previously (Degen et al., 2025, 2026; Gonzaga-Saavedra et al., 2026). After thawing, embryos (20 per stage per replicate) were resuspended in 700 µl RIPA buffer + protease inhibitor cocktail (Sigma P8340) (RIPA: 150 mM NaCl, 1% Igepal-CA630, 0.5% Sodium Deoxycholate, 0.1% SDS, 50 mM Tris pH7.4). Embryos were sonicated 4x 15 seconds at 20% output (Branson Sonifier 450, ¼-inch microtip horn) to shear chromatin. The sonication product for each stage (sheared chromatin in 700 µl RIPA buffer + protease inhibitors) was split in half into Eppendorf DNA Lo-Bind tubes (Eppendorf 022431021) and aliquots were brought up to 650 µl final volume with fresh RIPA buffer + protease inhibitors. The aliquots were incubated with primary antibodies (1 µg each per ChIP) overnight at 4°C on a nutator. Total H3 (Mouse monoclonal 1B1B2, Cell Signaling Technology 14269) and H3K27me3 (Rabbit Polyclonal, Diagenode C15410195) was ChIPped from stage 9, stage 11, and stage 12 samples for each replicate.

After the overnight incubation, 15 µl of Protein G Dynabeads (Fisher 10004D) pre-blocked overnight in PBS + 5% Bovine Serum Albumin + 0.2 mg/ml Yeast tRNA (Sigma 10109223001) was added and the samples were incubated at 4°C for an additional hour. After immobilization on a magnetic separation stand, bead-antibody complexes were washed 2x with 10 mM Tris pH 7.5 + 5 mM MgCl_2_. A tagmentation reaction was performed on the bead-antibody complexes in 20 µl of 1x Illumina Tn5 reaction buffer and 1 µl of Tn5 transposase (Illumina 20034210), incubating at 37°C with 1000 RPM shaking on an Eppendorf Thermomixer. 2x 5-minute washes in ice-cold Wash Buffer I (20 mM Tris-HCl pH 8.0, 0.1% SDS, 1% Triton X-100, 2mM EDTA, 150 mM HCl) were performed on the bead-antibody complexes on a nutator at room temperature, followed by 2x 5 minute washes in ice-cold Wash Buffer II (Wash Buffer I with 500 mM NaCl), and 1 x 5 minute wash in TE (10 mM Tris, pH 8.0, 1 mM EDTA). DNA elution and crosslink reversal was performed by incubating the bead-antibody complexes in 50 µl of Elution Buffer + Proteinase K (50 mM Tris pH8.0, 10 mM EDTA, 1% SDS + 1 mg/ml Proteinase K) shaking at 1200 RPM at 65°C. A Zymo ChIP DNA Clean and Concentrator kit (D5205) was used for DNA clean-up (following manufacturer’s instructions and eluting in 22.5 µl).

PCR was performed to amplify and barcode libraries. For PCR, 22.5 µl of ChIP DNA was combined with 2.5 µl of 25 mM Dual-unique barcoded primers and 25 µl of NEBNext 2x Hi-Fi PCR Mastermix (New England Biolabs M0541), with primers as defined previously (Buenrostro et al., 2013; Soluri et al., 2020). A QPCR side reaction was performed to determine the number of total PCR cycles for the main reaction by combining a 5 µl aliquot of the initial PCR solution with 5 µl of NEBNext Hi-Fi PCR Mastermix and 5 µl of SYBR-Primer mix (1.8x SYBR Green I (Invitrogen S7563) + 1.25 mM Dual-unique barcoded primer mix). The QPCR side reaction was set to one 5 minute cycle at 72°C and 30 seconds at 98°C and 25 cycles of 10 seconds at 98°C, 30 seconds at 63°C, and 1 minute at 72°C. At the end of each extension stage in the QPCR reaction, an Applied Biosystems Quantstudio 3 QPCR system imaged SYBR Green fluorescence. Dividing the plateau QPCR fluorescence value by 3 and rounding up to the nearest whole cycle number yielded the number of additional PCR cycles for the main PCR reaction. The main PCR was then run with equivalent cycling settings for the calculated number of additional cycles. 1.8x Ampure (Beckman Coulter A63880) was then used to clean-up the amplified libraries, with elution in 15 µl Qiagen Buffer EB. Qubit 1x dsDNA High Sensitivity fluorometry (Fisher Q33230) was used to measure library concentrations and TapeStation D5000 High Sensitivity reagents (Agilent 5067-5592) were used to measure fragment sizes. Equimolar concentrations of fragments sized between 100 and 700 bp were pooled together. An Illumina NovaSeq (Admera Health) was used to produce 150 bp paired-end reads.

### QUANTIFICATION AND STATISTICAL ANALYSIS

#### Sequencing data analysis

##### External datasets

Bigwigs from GEO: GSE299311 (Gonzaga-Saavedra et al., 2026) and GEO: GSE210236 (Bonnet et al., 2022) were used for analysis as described below. In Fig. S2B, H3K27me1, H3K27me3, E(z), and Pho data from GEO: GSE299311 (Gonzaga-Saavedra et al., 2026) is plotted, along with H3K27me2 data from GEO: GSE327938 (Degen et al., 2026). H3K27me3 measurements at CpG islands from Højfeldt et al. (Højfeldt et al., 2018) were used for comparison with the output of the mouse variation of the model.

##### Sequencing read mapping

As described previously (Degen et al., 2026), sequencing reads were trimmed using FastP (0.11.5, (Chen, 2023)) and mapped to the *Drosophila melanogaster* reference genome assembly (dm6) using bowtie2 (2.4.1, (Langmead & Salzberg, 2012)). In mapping, default parameters were used, with the exception of using the-x 2000 parameter to omit paired-end reads to 2000 bp lengths. Picard MarkDuplicates (2.21.4, https://broadinstitute.github.io/picard/) was used to mark PCR duplicates.

##### Creation of bigwigs

For our H3K27me3 ChIP-seq measurements at NC14 + 2h, + 4h, and + 6h,.bam files were imported to R using the GenomicAlignments package (1.34.0, (Lawrence et al., 2013)) keeping only properly-paired nonsecondary mappings with map quality greater than 10 and excluding duplicates. This made a GenomicRanges object. The average read coverage over 10 basepair windows was then calculated using the binnedAverage() function in the GenomicRanges package (1.50.1, (Lawrence et al., 2013)). Average coverage scores were normalized to counts per million and bigwigs were exported using the rtracklayer package (1.58.0, (Lawrence et al., 2009)).

For the analyses of H3K27me3 at NC14 + 15’, + 35’ and + 60’, bigwigs were downloaded from GEO: GSE299311 (Gonzaga-Saavedra et al., 2026). For the analysis of H3K27me3 in wild-type control and Sce[I48A] embryos, as well as the H3K27me2/3 analysis in 24h embryos, bigwigs were downloaded from GEO: GSE210236 (Bonnet et al., 2022). For each dataset, the average reads per 10 bp window was calculated.

##### Isolated PRE analysis

As described previously (Degen et al., 2026), E(z) ChIP-seq peaks (from GEO: GSE299311, (Gonzaga-Saavedra et al., 2026)) were chosen for analysis if they overlap with H3K27me3 in NC14, if they fall into the top 90% of E(z) peaks in terms of max signal, and if they are more than 27 kilobases (or 150 nucleosomes, at 180 bp/nucleosome) away from neighboring E(z) peaks. The selected E(z) peaks are referred to as “isolated PREs.”

##### Spreading rate calculations

To determine the *in vivo* H3K27me3 spreading rate, exponentials were fit to the H3K27me3 ChIP-seq measurements at isolated E(z) peaks across the 8th to 150th proximal nucleosomes at NC14 + 35’, NC14 + 60’, NC14 + 120’, NC14 + 240’, and NC14 + 360’. In the fitting process, H3K27me3 signal distal to the first proximal boundary element associated with each E(z) peak was excluded, thereby reducing the influence of topological constraints on H3K27me3 spreading. The decay lengths of the exponentials were then plotted against the time points of the measurements. For the NC14 + 15’ time point, the decay length was set to 0, as at this early time no H3K27me3 has yet accumulated. A logarithmic curve was then fit to the plotted decay lengths, and the derivative of the curve calculated to determine the *in vivo* H3K27me3 spreading rate.

To determine the modeled H3K27me3 spreading rate, exponentials were fit to the predicted H3K27me3 distributions across the 8th to 150th nucleosomes flanking the modeled PRE at NC14 + 35’, NC14 + 60’, NC14 + 120’, NC14 + 240’, and NC14 + 360’. To match our treatment of the *in vivo* data, an exponential was not fit to the predicted H3K27me3 at NC14 + 15’. The decay lengths of the exponential fits to the modeled H3K27me3 were then plotted against the time points of the measurements. For the NC14 + 15’ time point, the modeled decay length was set to 0. A logarithmic curve was then fit to the plotted decay lengths, and the derivative of the curve calculated to determine the modeled H3K27me3 spreading rate.

#### Modeling

##### Initial conditions

In a single nucleus, for each chromosome there are two homologs, and therefore at a given nucleosome position in the genome, there are four H3K27 positions that can be possibly methylated by E(z). Given the average diameter of an NC14 nucleus (∼3 um, based on prior live imaging measurements (Gonzaga-Saavedra et al., 2026)), we calculated the nuclear volume assuming the nucleus is a sphere, and then calculated the maximal concentration of H3K27 methylation that could accumulate at a nucleosome position in the genome: 0.06 nM. To set the initial conditions of the model, we assumed 50% of the H3K27 positions at the modeled 300-nucleosome array are unmethylated, 25% the positions contain me1, and 25% of the positions contain me2, given recent computational and experimental work suggesting these H3K27 states exist at the start of NC14 (Degen et al., 2026). The model’s initial conditions lack H3K27me3, as experimental measurements indicate the modification is absent from chromosomes at the start of NC14 (Gonzaga-Saavedra et al., 2026; Li et al., 2014; Reinig et al., 2020), supported by computational modeling (Degen et al., 2026; Lundkvist et al., 2023). We specify in our model that the initial distributions of H3K27me1 and-me2 are uniform across the nucleosome array, given the broad distribution of H3K27me2 measured in cleavage-stage *Drosophila* embryos (Degen et al., 2026). Given these considerations, at the start of NC14, we model the chromatin fiber as having uniform 0.03 nM me0, 0.015 nM me1, and 0.015 nM me2.

We evaluated how much the model’s outputs depend on its initial conditions by varying the starting state of the modeled nucleosome array. Increasing the level of H3K27me2 on the array at the start of NC14 does not significantly impact the model’s prediction of the establishment of an H3K27me3 domain (Fig. SM1 A-F).

**Figure Supplemental Methods 1.**
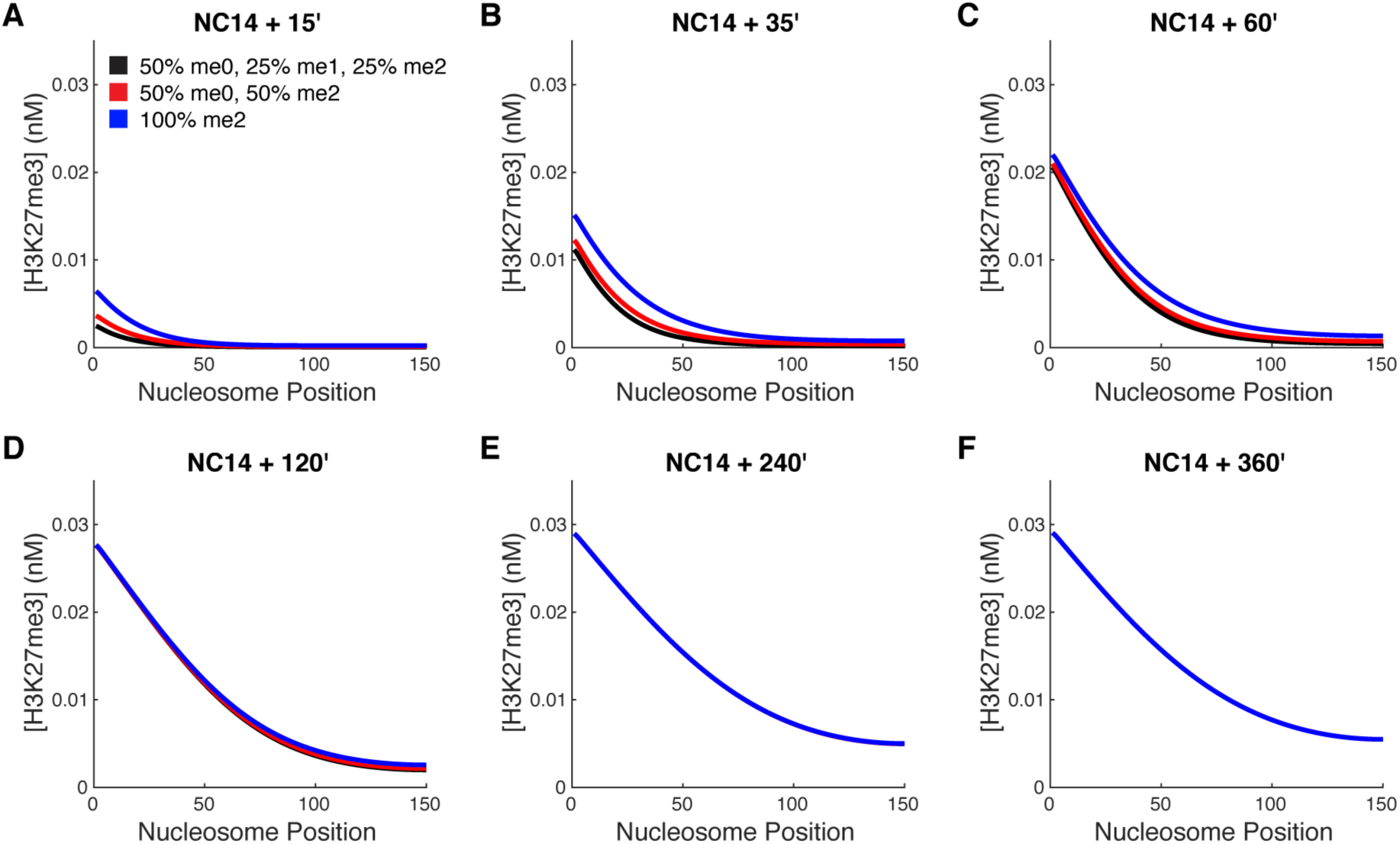
Testing initial conditions. Shown are simulated H3K27me3 distributions upon varying the model’s initial conditions. Nucleosome position = 0 marks the position of the PRE on the modeled chromatin fiber. The legend specifying the initial conditions in (A) applies to all panels. The legend indicates the percent of the H3K27 positions in the nucleosome array that start NC14 with me0/1/2/3, with 100% H3K27 corresponding to 0.06 nM. All initial conditions were uniform across the modeled chromatin fiber. All simulations used the parameter values defined in Table SM1.

For all simulations, E(z) was absent from the nucleosome array at the start of NC14 (*t* = 0), given the enzyme is evicted from nuclei during mitosis (Gonzaga-Saavedra et al., 2026).

##### Numerical simulations

The reaction-diffusion system was solved numerically in MATLAB (R2022a) with the pdepe function using no-flux Neumann boundary conditions. All simulations were run for a 300 nucleosome array with a PRE positioned at the 150th nucleosome (*x* = 0). *t* = 0 corresponded to the start of NC14 for all simulations.

##### Allostery decision

Recent work has predicted that allosteric stimulation of E(z) is key for the rapid establishment of H3K27me3 in the *Drosophila* embryo (Degen et al., 2026). While the model presented here does not incorporate allostery explicitly, it remains compatible with a role for Esc in the H3K27me3 establishment process. H3K27me2 is maintained on chromatin through early development, and could act to effectively raise the catalytic efficiency of E(z) (Degen et al., 2026). The model’s elevated E(z) efficiency compared with the *in vitro* efficiency estimate may reflect the stimulation of E(z) by Esc interacting with dimethylation in the early embryo (Sneeringer et al., 2010). Due to its high catalytic efficiency setting, the model successfully predicts the observed H3K27me3 dynamics without needing to nonlinearly relate E(z) reaction rates to substrate concentrations. As adding nonlinearity to our model would hinder our ability to equate *R* to the efficiency of the enzyme, in this work we focused on the simpler form of the reaction-diffusion framework. The model therefore provides a platform for quantifying how eliminating the ability of Esc to allosterically stimulate E(z) changes the *in vivo* catalytic efficiency of the enzyme.

#### Parameter summary

**Table Supplemental Methods 1.** Parameter values. Shown are the parameter values used in the reaction-diffusion system to model H3K27me3 in wild-type *Drosophila* embryos.

| Parameter | Value | Source |
| --- | --- | --- |
| Baseline E(z) catalytic efficiency, $R$ | $9.3 \times 10^{-5} \text{ nM}^{-1} \text{ min}^{-1}$ | This paper |
| Rate constant for E(z) nucleation at the PRE, $k_{on}$ | $6 \text{ min}^{-1}$ | This paper |
| Rate constant for background E(z) association with a nucleosome, $k_b$ | $0.01 \text{ min}^{-1}$ | This paper |
| Rate constant for E(z) dissociation from the random walk, $k_{off}$ | $0.02 \text{ min}^{-1}$ | This paper |
| Nuclear E(z) concentration, $E_z$ | 60 nM | Degen et al., 2026 (Degen et al., 2026) |
| 1D diffusion coefficient for $E(z)$, $D$ | 55 nucleosomes<sup>2</sup> min<sup>-1</sup> | This paper, calculated from measurements in Gonzaga-Saavedra et al., 2026 (Gonzaga-Saavedra et al., 2026) |
| Rate constant for Utx-independent demethylation, $d_H$ | 1/90 min<sup>-1</sup> + 1/90 min<sup>-1</sup> | This paper, based on Deal et al., 2010 and Hartenstein & Campos-Ortega, 1985 (Deal et al., 2010; Hartenstein & Campos-Ortega, 1985) |
| Rate constant for Utx-dependent demethylation, $d_{UTX}$ | 3.125 x 10<sup>-5</sup> min<sup>-1</sup> | This paper, calculated from measurements in Zheng et al., 2012 (Y. Zheng et al., 2012) |

##### Definition of *d_H_*

*d_H_* is the rate constant for the demethylation of H3K27 due to nucleosome turnover and the incorporation of unmodified histones to chromatin at DNA replication. This parameter is calculated from the average nucleosome lifetime at PcG protein-bound sites in *Drosophila* cells (90 minutes, (Deal et al., 2010)), as well as the frequency of DNA replication from mitosis 13 to 6 hours post-mitosis 13 (on average every 90 minutes, (Hartenstein & Campos-Ortega, 1985)). In the model, we account for the effects of histone modification dilution during genome replication continuously, given the mitoses of cells within the embryo are asynchronous at these stages, and the ChIP-seq measurements that we benchmark our model against are an average of all cellular states in the embryo. Further, the genome does not replicate instantaneously within each cell, and therefore modeling replication-dependent histone dilution continuously over time is more appropriate than modeling dilution at a single time point. With these considerations, *d_H_* = 1/90 min^-1^ + 1/90 min^-1^.

##### Definition of *d_UTX_*

*d_UTX_* is the rate constant for demethylation of H3K27 due to activity of the H3K27 demethylase Utx. As it has been reported Utx only acts on H3K27me2/3 (Smith et al., 2008), *d_UTX_* only mediates the me3-me2 and me2-me1 transitions in the model. We calculated *d_UTX_* by averaging *in vivo* measurements of the effective rate constants for me3-me2 and me2-me1 transitions made in mammalian cell culture (Y. Zheng et al., 2012). As these measurements were not made in *Drosophila* cells, we evaluate the dependency of the model on *d_UTX_* in the methods.

##### Definition of *k_off_*

*k_off_* is the rate constant for E(z) dissociation from its random walk along chromatin. We estimate that on average E(z) spends 50 minutes engaging with the nucleosomes an a Polycomb domain, as we calculated the diffusion coefficient for the 1D random walk of E(z) along chromatin from H3K27me3 domain widths produced after an average of 50 minutes post-mitosis 13. Therefore, our model uses *k_off_*= 1/50 min^-1^ = 0.02 min^-1^.

##### Definition of *D*

*D* is the diffusion coefficient governing the 1D random walk of E(z) along a nucleosome array containing a nucleation site. As discussed in the main text, *D* was calculated using Eq. 3 given the average H3K27me3 domain radius (the displacement of E(z) from a PRE after an average of 50 minutes following mitosis 13) is 13.3 kilobases.

##### Definition of *E_z_*

*E_z_* is the nuclear E(z) concentration at NC14, as calculated previously (Degen et al., 2026).

##### Wild-type parameter fitting

As discussed in the main text, we determined values for *k_on_*, *k_b_*, and *R* by performing parameter sweeps and evaluating how well the model predicted *in vivo* measurements at each parameter setting. To fit *k_on_* and *k_b_*, we swept across *k_on_* = 1 min^-1^ to 10 min^-1^ in steps of 1 min^-1^, and *k_b_* = 0 min^-1^ to 0.1 min^-1^ in steps of 0.0025 min^-1^. All other parameters were held at their baseline values specified in Table SM1, except for *R*, which was held at an arbitrary value of 1 as it does not influence the distribution of E(z) on chromatin. At each *k_on_* and *k_b_* combination, we calculated the difference between the estimated nuclear E(z) concentration (60 nM, (Degen et al., 2026)) and the average modeled E(z) concentration at the 300 nucleosome array over NC14. Minimizing the squared residual indicated the best-fit parameter combination is *k_on_* = 6 min^-1^, *k_b_*= 0.01 min^-1^ (Fig. 3C).

To determine the best-fit *R* value, we swept from *R* = 10^-6^ (nM^-1^ min^-1^) to 10^-3^ (nM^-1^ min^-1^) in steps of 10^-6^ (nM^-1^ min^-1^), and calculated the sum of squared residuals comparing the predicted H3K27me3 decay length at each time point (NC14 +35’, +60’, +120’, +240’, +360’) to the decay lengths of the exponential fits to the mean H3K27me3 measurements at each time point (Fig. 2B & C). All other parameters were held at their baseline values specified in Table SM1. Predicted decay lengths were determined by fitting exponentials to the model’s H3K27me3 predictions from *x* = 8 to 150 to mimic the range over which exponentials were fit to our isolated PRE H3K27me3 data (the first seven nucleosomes proximal to the PRE were excluded). The NC14+15’ time point was excluded from the fitting process, as there is no detectable *in vivo* H3K27me3 at that time, and therefore calculating an interpretable decay length for the H3K27me3 measurement is impossible. Minimizing the sum of squared residuals indicated that *R* = 9.3 x 10^-5^ nM^-1^ min^-1^ allows the model to best predict the *in vivo* H3K27me3 spreading data (Fig. 3D, Fig. S2 A).

##### Testing parameter dependency

We evaluated the dependency of the model on its parameter values by performing parameter sweeps, varying each parameter individually while holding the others at their baseline values specified in Table SM1. For each parameter combination, we evaluated the model’s H3K27me3 prediction at NC14 + 360’. We find altering *k_on_* primarily affects H3K27me3 levels at the PRE (*x* = 0) (Fig. SM2 A), while changing *k_b_*primarily impacts H3K27me3 concentrations farther from the PRE (Fig. SM2 B). Increasing *k_on_* and *k_b_* allow more E(z) to accumulate on the nucleosome array, therefore increasing predicted H3K27me3 levels (Fig. SM2 A & B). Changing *R* and *E_z_* have similar effects; both parameters increase the magnitude of the predicted H3K27me3 distribution across the nucleosome array, with larger effects at the PRE (Fig. SM2 C & D). These parameters increase the rate of H3K27 methylation production. Alterations to *k_off_* change the spread of the modeled H3K27me3 distribution (Fig. SM2 E). Decreasing *k_off_* increases the residence time of E(z) on the nucleosome array, thereby allowing the enzyme to diffuse farther out from the PRE, and allowing more H3K27me3 to accumulate at each nucleosome (Fig. SM2 E). Changing *D* has a dramatic effect on the predicted spread of H3K27me3, with low *D* values producing a steep H3K27me3 boundary, while high *D* values allow E(z) to diffuse farther from the PRE, thereby producing a shallower H3K27me3 distribution (Fig. SM2 F). The magnitude of H3K27me3 close to the PRE inversely correlates with *D*, while the magnitude of H3K27me3 far from the PRE positively correlates with *D* (Fig. SM2 F). This observation highlights how the longer E(z) stays at a nucleosome, the more H3K27 methylation is catalyzed at that nucleosome. Changing *d_H_* changes the magnitude of the predicted H3K27me3 almost uniformly across the domain, with H3K27me3 levels inversely correlating with the magnitude of *d_H_*(Fig. SM2 G). Notably, across the range of tested *d_UTX_* values, no change to this parameter altered the predicted H3K27me3 distribution at NC14 + 360’ (Fig. SM2 H). Neither setting *d_UTX_* = 0 nor increasing *d_UTX_* to over twice its baseline value impacts the model’s H3K27me3 prediction at NC14 + 360’.

**Figure Supplemental Methods 2.**
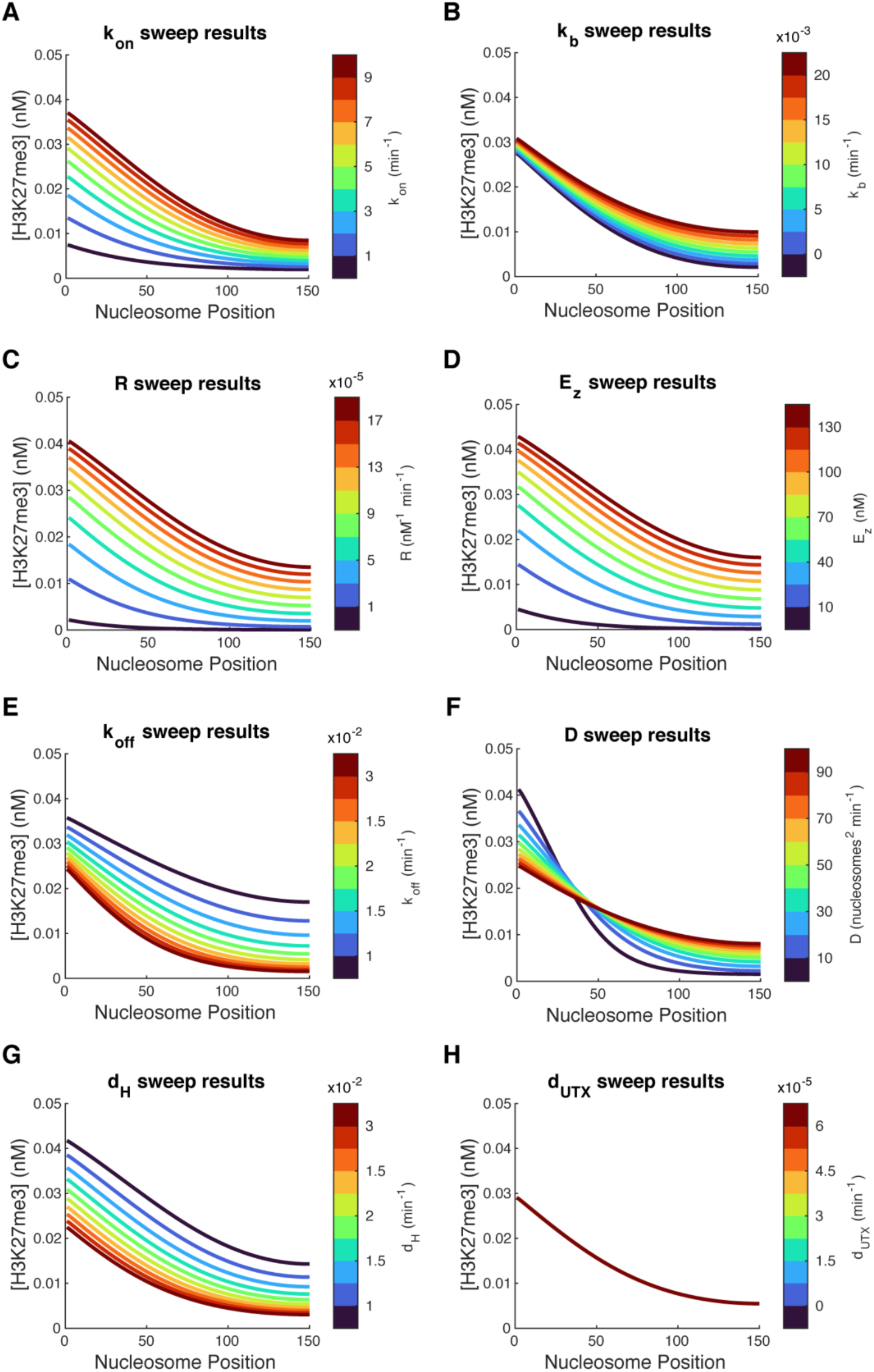
Testing parameter dependency. Shown are simulated H3K27me3 levels at NC14 + 360’ predicted by the model upon varying the specified parameters. Nucleosome position = 0 marks the position of the PRE on the modeled chromatin fiber. In A-H, all unspecified parameters were held at their baseline values presented in Table SM1.

##### Equilibrium simulation

To simulate H3K27me2/3 in 24h old embryos, we added a time dependence to *d_H_*, reducing this parameter from 2/90 min^-1^ to 1/90 min^-1^ after NC14 + 360’, as the cell cycle halts for most cells in late-stage *Drosophila* embryos, eliminating the contribution of DNA replication to histone turnover. All other parameters were held at their baseline values specified in Table SM1. To generate predictions for the 24h stage, we simulated from *t* = 0 (the start of NC14) to *t* = 22h, as NC14 begins ∼2 hours into development.

##### Sce[I48A] parameter fitting

We performed parameter sweeps to determine how changes to the model’s parameters allow it to recapitulate H3K27me3 measurements in Sce[I48A] mutants. We swept across *k_on_* (1 to 10 min^-1^ in steps of 1 min^-1^), *k_b_* (0 to 0.02 min^-1^ in steps of 2.5 x 10^-3^ min^-1^), *R* (1 x 10^-5^ to 2 x 10^-4^ nM^-1^ min^-1^ in steps of 2 x 10^-5^ nM^-1^ min^-1^), and *D* (5 to 70 nucleosomes^2^ min^-1^ in steps of 5 nucleosomes^2^ min^-1^), and at each parameter combination, determined how the model’s H3K27me3 prediction at NC14 + 120’ compared to the wild-type model’s H3K27me3 prediction at NC14 = 120’ (using the parameters specified in Table SM1). We chose to evaluate the model’s output at NC14 + 120’, as the 0-6h old embryo data we were attempting to simulate primarily captures DNA from post-cleavage stage embryos in the time range (NC14 + 0’ to NC14 + 240’), given the higher cell content at these stages. The average state in the 0-6h data therefore likely corresponds to NC14 + 120’. At each parameter combination, to the model’s H3K27me3 prediction at NC14 + 120’ we fit an exponential,

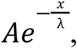

and determined how well the parameter “mutation” predicted the relative changes in A and λ seen in the exponential fits to the wild-type and Sce[I48A] measurements. Decreasing *k_on_*, *k_b_*, *R*, and *D* from their wild-type settings produced the most accurate predictions of the drop in H3K27me3 magnitude and spread produced by the Sce[I48A] mutation (Fig. S3).

The top 1% of parameter sets that best predicted the H3K27me3 distribution in Sce[I48A] contain the following averages: *k_on_* = 4 min^-1^, *k_b_* = 0.007 min^-1^, *R* = 6 x 10^-5^ nM^-1^ min^-1^, and *D* = 19 nucleosomes^2^ min^-1^. Compared to the “wild-type” parameter set (*k_on_* = 6 min^-1^, *k_b_* = 0.01 min^-1^, *R* = 9.3 x 10^-5^ nM^-1^ min^-1^, and *D* = 55 nucleosomes^2^ min^-1^), these mutant parameter values mark a 33% decrease in *k_on_*, 30% decrease in *k_b_*, 35% decrease in *R*, and 65% decrease in *D*. For each parameter, the majority of values that best predict the Sce[I48A] data fall below the wild-type setting (Fig. S3), indicating these parameters must be simultaneously decreased for accurate predictions.

We also evaluated the results of parameter combinations when *D* = 20 nucleosomes^2^ min^-1^, as this value approximates the mean *D* in the top 1% parameter sets (Fig. S3), and is therefore associated with a substantial number of successful parameter sets (Fig. SM3 A-J). At *R* values close to the mean value of the top 1% parameter sets (*R* = 6 x 10^-5^ nM^-1^ min^-1^), when *D* = 20 nucleosomes^2^ min^-1^, the model best predicts the Sce[I48A] data when *k_on_* = ∼3-4 min^-1^ (Fig. SM3 C & D), values notably lower than the model’s baseline *k_on_* value of 6 min^-1^. At moderately low *R* values, when *D* = 20 nucleosomes^2^ min^-1^, the model depends less on its precise *k_b_* setting to make accurate predictions, however, it does have a slight preference for lower *k_b_* settings (Fig. SM3 C & D). Together, these results suggest that simultaneously lowering *k_on_*, *k_b_*, *R*, and *D* in general most appropriately shifts the model’s output to capture the H3K27me3 distribution in Sce[I48A] embryos. However, at very low *R* values, the model’s ability to predict the Sce[I48A] data significantly suffers (Fig. SM3 A), indicating the model requires a fairly precise definition of *R*.

**Figure Supplemental Methods 3.**
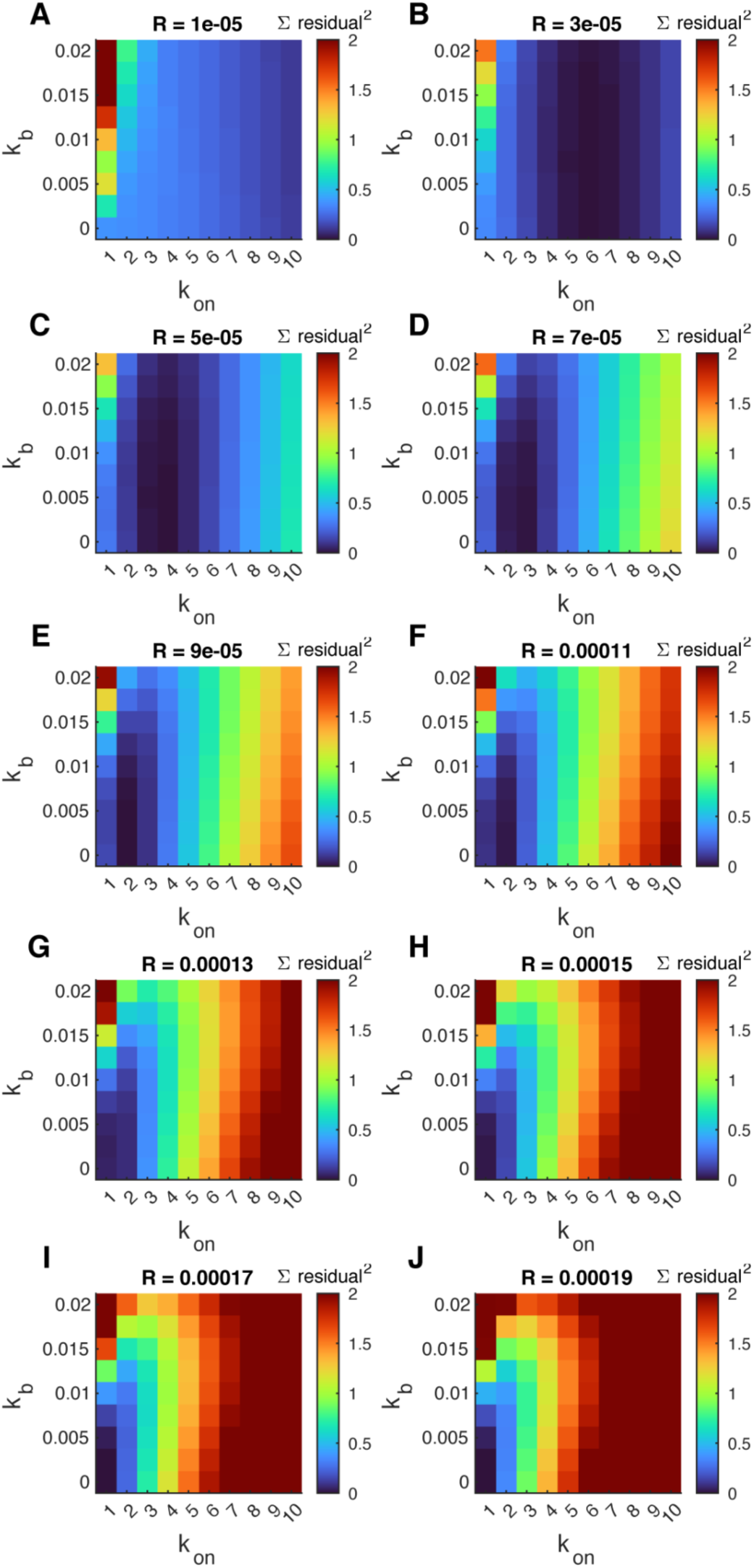
Sce[I48A] parameter sweep results at D = 20. Heatmaps show the sum of squared residuals calculated by comparing the modeled λ_Sce_/λ_WT_ and A_Sce_/A_WT_ to the ratios calculated from the exponential fits to Sce[I48A] and wild-type control H3K27me3 measurements from Bonnet et al (Bonnet et al., 2022). In A-J all axes have units of min^-1^. *R* values have units of nM^-1^ min^-1^. Sweep results when *D* = 20 are shown, as at this value, a significant number of *R*, *k_on_*, and *k_b_* combinations produce good predictions.

##### Mouse model variation

**Table Supplemental Methods 2.** Parameter values for the mouse model. Shown are the parameter values used in the reaction-diffusion system to model H3K27me3 in mouse embryos.

| Parameter | Value | Source |
| --- | --- | --- |
| Baseline E(z)/EZH2 catalytic efficiency, $R$ | $5.7 \times 10^{-6} \text{ nM}^{-1} \text{ min}^{-1}$ | Sneeringer et al., 2010 (Sneeringer et al., 2010) |
| Rate constant for E(z)/EZH2 nucleation, $k_{on}$ | $6 \text{ min}^{-1}$ | This paper |
| Rate constant for background E(z)/EZH2 association with a nucleosome, $k_b$ | $0.01 \text{ min}^{-1}$ | This paper |
| Rate constant for E(z)/EZH2 dissociation from the random walk, $k_{off}$ | $0.02 \text{ min}^{-1}$ | This paper |
| Nuclear E(z)/EZH2 concentration, $E_z$ | 10 nM | Based on the estimation for mammalian cells from Zheng et al., 2012 (Y. Zheng et al., 2012) |
| 1D diffusion coefficient for E(z)/EZH2, $D$ | $55 \text{ nucleosomes}^2 \text{ min}^{-1}$ | This paper, calculated from measurements in Gonzaga-Saavedra et al., 2026 (Gonzaga-Saavedra et al., 2026) |
| Rate constant for enzyme-independent demethylation, $d_H$ | $0.0017 \text{ min}^{-1}$ | Combining cell cycle and histone turnover measurements in mESCs (Deaton et al., 2016; Waisman et al., 2019) |
| Rate constant for enzyme-dependent me1 demethylation, $d_{10}$ | $1.4583 \times 10^{-4} \text{ min}^{-1}$ | Zheng et al., 2012 (Y. Zheng et al., 2012) |
| Rate constant for enzyme-dependent me2 demethylation, $d_{21}$ | $4.8611 \times 10^{-5} \text{ min}^{-1}$ | Zheng et al., 2012 (Y. Zheng et al., 2012) |
| Rate constant for enzyme-dependent me3 demethylation, $d_{31}$ | $1.3889 \times 10^{-5} \text{ min}^{-1}$ | Zheng et al., 2012 (Y. Zheng et al., 2012) |

##### Initial conditions

For the mouse model simulations, we used an initial condition of uniform 0.06 nM H3K27me1, as this modification was maintained on chromatin while H3K27me2/3 was largely depleted upon treatment of mESCs with an EZH2 inhibitor (Højfeldt et al., 2018). Therefore, in the time course data we intended to simulate (Højfeldt et al., 2018), chromatin started with H3K27me1 at *t* = 0. For comparison with the mouse model, we also ran a simulation with the *Drosophila* model using an initial condition of uniform 0.06 nM H3K27me1 (results shown in Fig. 4D).

##### Demethylation implementation

There is evidence in mammals that H3K27me1,-me2, and-me3 can all be actively demethylated (Hong et al., 2007). We incorporated reverse transitions between H3K27 methylation states based on effective rate constants determined for mammalian cells (*d_10_*, *d_21_*, and *d_32_* reported in Table SM2) (Y. Zheng et al., 2012).

##### Comparison to *in vivo* data

To compare the model’s output to *in vivo* measurements of H3K27me3 at CGIs, we integrated the predicted H3K27me3 levels along the modeled chromatin fiber at each time point, and normalized the resultant values by the maximum over the simulated 24 hours. To compare these estimates to *in vivo* data, we drew upon prior ChIP measurements of H3K27me3 at CGIs (Højfeldt et al., 2018). We downloaded the source data for Fig. 3 of Højfeldt et al. and found the averages and standard deviations of the H3K27me3 measurements at Dclk1 and Pmp22 loci over time. We then normalized these values by the maximum H3K27me3 average over the 24 hours post-inhibitor washout.

## Author contributions

**E.A.D.:** original draft preparation; review and editing; conceptualization; methodology; investigation; software; formal analysis; data curation; visualization. **S.A.B.:** supervision and administration; funding acquisition; review and editing; investigation.

## Acknowledgments

We are grateful to R. Braun for her feedback on this work and for her comments on the manuscript. We also thank J. Marko for his feedback on the project. We thank the Bloomington Drosophila Stock Center and Flybase for providing essential resources to the *Drosophila* community. E.A.D. was supported by the Cellular and Molecular Basis of Disease training program (T32 GM008061), is supported by an NSF GRFP fellowship (DGE-2234667), and is a Data Science Fellow at the Northwestern Institute on Complex Systems. Experiments were supported by the Alumnae of Northwestern University and the National Institutes of Health grant R01 HD101563 to S.A.B.. S.A.B. is a Pew Scholar in the Biomedical Sciences, supported by the Pew Charitable Trusts.

